# Integrative transcriptomic and phosphoproteomic analysis reveals key components of SnRK1 signaling network in rice

**DOI:** 10.1101/2025.07.22.666209

**Authors:** Maria C. Faria-Bates, Chandan Maurya, K Muhammed Jamsheer, Vibha Srivastava

## Abstract

SnRK1 is an evolutionarily conserved protein kinase belonging to SNF1/AMPK family of protein kinases that is central to adjusting growth in response to the energy status. Numerous studies have shown adaptive and developmental roles of SnRK1, but the understanding of SnRK1 signaling network in monocots is limited. Using CRISPR/Cas9 mutagenesis to target the functional kinase subunits in rice, we carried out a comprehensive phenotypic, transcriptomic, proteomic, and phosphoproteomic analyses of rice *snrk1* mutants displaying growth defects under normal and starvation conditions. These analyses revealed the role of SnRK1 signaling in controlling growth and stress-related processes in both energy-sufficient and energy-limited conditions and pointed to sub-functionalization of SnRK1 kinase subunit genes. In addition to the classical protein targets of SnRK1, phosphoproteomics revealed novel targets including the key components of intracellular membrane trafficking, ethylene signaling, and ion transport. The upregulation of stress-related processes and suppression of growth-related processes in *snrk1* mutants correlated with their phenotypic defects. Overall, this study highlights the dual role of SnRK1 as promoter of growth under favorable conditions and critical regulator of adaptive response under stress conditions.

## INTRODUCTION

Sucrose Non-Fermenting Related Kinase 1 (SnRK1) plays a crucial role in coordinating growth and cellular metabolism according to energy status through phosphorylation of its target proteins in Ser/Thr sites. SnRK1 is the plant ortholog of the evolutionarily conserved SNF1/AMPK protein kinase family and is activated by starvation and other stress conditions. The hetero-trimeric SnRK1 complex contains three subunits: the catalytic α subunit, and the regulatory β and βγ subunits (Broeckx et al., 2016; Jamsheer et al., 2021; Polge & Thomas, 2006). To maintain cellular energy homeostasis and ensure plant’s survival, SnRK1 promotes catabolism and suppresses anabolism under stress (Peixoto & Baena-Gonzalez 2022; Cho Y-H et al., 2012; Pedrotti et al., 2018; Wang et al., 2021), and possibly also in the normal or energy-sufficient conditions (Henninger et al., 2022; Wang et al., 2021). Upon activation, SnRK1 triggers signaling events that lead to the repression of anabolic processes such as cell wall formation, protein translation, and ribosome biogenesis and induction of catabolic processes, *e.g*., carbohydrate, lipid, and amino acid catabolism (Baena-Gonzalez et al., 2007; Henninger et al., 2022; Lu et al., 2007; Nukarinen et al., 2016; Wang et al., 2021).

It is now well-established that SnRK1 controls metabolic processes through phosphorylation of key metabolic enzymes that affects their stability and/or activity. Some of the known SnRK1 targets are the sucrose-phosphate synthases (SPS), fructose-2,6-bisphosphatase (F2KP), carbonic anhydrase (CA), pyruvate kinase (PK), sucrose synthase (SUS), inositol polyphosphate kinase 2 beta (IPK2β), nitrate reductase (NR), and diacylglycerol acyltransferase 1 (DGAT1) (Caldo et al., 2018; Cho et al., 2016; Kulma et al., 2004; Luo et al., 2020; Song et al., 2019; Sugden et al., 1999; Yang et al., 2018). By phosphorylating these proteins, SnRK1 regulates carbohydrate, inositol, nitrogen, and lipid metabolism. Additionally, SnRK1 regulates numerous anabolic processes through inhibition of Target of Rapamycin Complex 1 (TORC1) signaling by phosphorylating TORC1 component, RAPTOR1 (Gwinn et al., 2008; Nukarinen et al., 2016). SnRK1 is also known to orchestrate signaling cascades by interacting with different protein kinases and phosphatases. Protein-Protein Interaction (PPI) networks in Arabidopsis have revealed the role of SnRK1 in defense pathways through its interaction with mitogen-activated protein kinase 6 (MAPK6) and other MAPK domain proteins and receptor-like kinase (RLK) family proteins (Carianopol et al., 2020; Cho et al., 2016; Jamsheer et al., 2021). Moreover, by stimulating jasmonic acid (JA) and salicylic acid (SA) signaling in rice, SnRK1 promotes broad-spectrum disease resistance against bacterial and fungal pathogens (Cao et al., 2024; Filipe et al., 2018). Arabidopsis SnRK1 interacts with proteins involved in biotic stress such as Phloem protein 2 A5 (PP2A5), nematode resistance proteins, HSPRO1 and HSPRO2, and directly phosphorylates non-expressor of PR genes (NPR1) to control plant immunity (Chen et al., 2025; Jamsheer et al., 2021). Thus, by integrating plant growth, development, and defense responses, SnRK1 acts as a signaling hub, triggering adaptive responses to promote plant survival in unfavorable conditions. Similarly, SnRK1 also regulates gene expression by phosphorylating transcription factors and chromatin-remodeling enzymes (Chan et al., 2017; Mair et al., 2015; Wang et al., 2021).

Our knowledge of the SnRK1 signaling network in plants is mostly based on the studies conducted on Arabidopsis. Therefore, our understanding of SnRK1 signaling in monocots is rather limited. Nonetheless, studies so far indicate a crucial adaptive and developmental role of SnRK1 in monocots such as rice and maize (Filipe et al., 2018; Li et al., 2020; Wang et al., 2021, Yang T et al., 2024). To map the comprehensive network of SnRK1 signaling and resolve its conservation and diversity across the plant lineage, more studies focusing on SnRK1 signaling networks in groups such as monocots are needed. This information is also important in the deeper understanding of SnRK1-dependent signaling network and its relationship with growth and adaptive responses in plants.

In this study, we focused on the three functional kinase subunits of SnRK1 to systematically analyze the SnRK1-dependent signaling network in rice. We investigated the role of the SnRK1 signaling in the growth and adaptation of rice seedlings to starvation through phenotypic characterization of rice SnRK1 mutants developed through CRISPR/Cas9 mutagenesis and their transcriptomic, proteomic, and phosphoproteomic analyses. We found that SnRK1 plays a major role in the normal growth and development, energy starvation, and defense against rice blast fungus (*Magnaporthe oryzae*). Through transcriptomic analysis, we identified the SnRK1-dependent gene networks involved in the developmental processes and adaptive responses to starvation. Through proteomics and phosphoproteomics, we identified protein networks regulated by SnRK1. Further, along with the classical conserved targets such as RAPTOR, we identified potentially novel phosphosites of the SnRK1 signaling network in rice. Together, these findings highlight the central role of SnRK1 as a hub for coordinating not only stress and defense responses but also energy-driven growth processes in rice and possibly other plant species.

## RESULTS

### SnRK1 is a key regulator of plant growth and defense in rice

In a previous study, we developed rice mutants of SnRK1 kinase subunits through CRISPR/Cas9 mutagenesis (Pathak et al., 2022). Two sets of *snrk1* mutants were developed based on sequence homology of the three functional paralogs of the SnRK1 kinase α-subunit genes (*OsSnRK1*α): *OsSnRK1*α*A* (Os05g0530500)*, OsSnRK1*α*B* (Os8g0484600), and *OsSnRK1*α*C* (Os03g0289100). Of these, *OsSnRK1*α*B* and *OsSnRK1*α*C* show high sequence homology (88.2% genomic and 98.02% protein sequence), while *OsSnRK1*α*A* is more divergent. Based on this, single-mutants of *OsSnRK1*α*A* and double-mutants of *OsSnRK1*α*BC* were developed using the *japonica* rice variety, Kitaake, and the characterized homozygous lines of *ossnrk1*α*a* and *ossnrk1*α*bc* were selected for this study. These mutants are referred to as *snrk1a* and *snrk1bc*, hereafter. Both mutants contain truncation in the kinase domain through incorporation of an early stop codon (**File S1**). Gene expression analysis by qPCR verified that the non-targeted paralog(s) of *OsSnRK1*α were not affected while the targeted paralog(s) were downregulated in each mutant (**Fig. S1**), indicating degradation of the nonsense transcript, a conserved gene regulatory mechanism in eukaryotes, and confirming the absence of collateral effects on the non-targeted paralogs.

To understand the role of SnRK1 in the seedling growth, the *snrk1a* and *snrk1bc* mutants were cultivated along with the wildtype (WT). Defective growth was observed in *snrk1*seedlings grown on half-strength MS media under 14 h photoperiod, representing normal condition (energy-sufficiency) as well as in seedlings exposed to 48 h of extended darkness inducing starvation (energy-deficiency). In normal condition, 9-d-old seedlings of *snrk1a* and *snrk1bc* mutants developed shorter length and lower biomass (shoot and root) compared to the wildtype (WT) (**Fig. 1a-b; Fig. S2a-d**). Under starvation, on the other hand, both *snrk1a* and *snrk1bc* seedlings appeared thinner than WT (**Fig. 1c**). Shoot and root length was not significantly different (**Fig. S2e-f**); however, reduced biomass was observed in *snrk1bc* seedlings **(Fig. 1d; Fig. S2g-h)**. Next, chlorophyll content of the seedlings under normal condition was not significantly different between the three genotypes (**Fig. 1e**), but under starvation, both *snrk1* mutants accumulated lower chlorophyll with *snrk1bc* seedlings showing a significant reduction (**Fig. 1f**). Most notable growth defects under starvation were observed in *snrk1bc* seedlings, *e.g.,* development of very thin seedlings, yellowing in the culm, and leaf rolling (**Fig. S3**). These observations indicate the role of SnRK1 signaling in rice seedling establishment and their response to starvation, as well as potential sub-functionalization of *OsSnRK1*α genes. The mature plants of *snrk1a* and *snrk1bc* mutants raised in the greenhouse condition also showed growth defects (**Fig. 1g-h**). Both *snrk1* mutants showed lower shoot and root biomass (**Fig. 1i-j**). Reduced fertility in *snrk1* mutants was indicated by lower number of seeds per panicle, while the weight of 100 grains was not significantly different (**Fig. 1k-l**). Here, *snrk1a* plants showed more severe reduction in biomass and fertility, indicating important roles for SnRK1 signaling in the vegetative and reproductive development of the plant. These results further confirm the functional diversification of *OsSnRK1*α genes.

**Figure 1:**
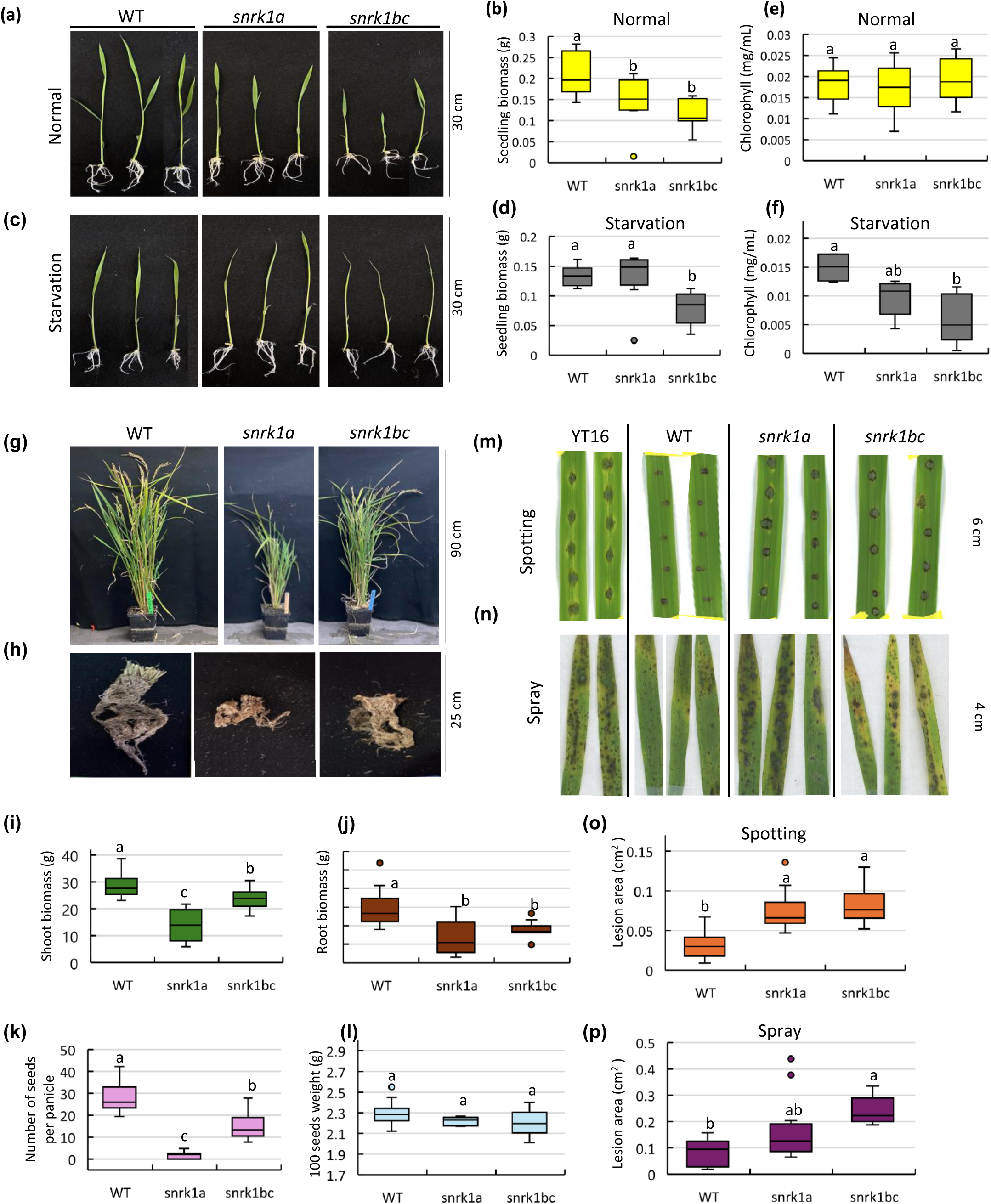
Phenotypic characterization of *snrk1* mutants. **(a-d)** Seedling phenotype and biomass of 9-d-old seedlings of wildtype Kitaake (WT), *snrk1a*, and *snrk1bc* grown on half-strength MS medium under 14 h photoperiod (normal condition) (a-b) or exposed to 48 h of extended darkness after 7 d of normal growth (starvation) (c-d). **(e–f)** Chlorophyll content in 9-d-old seedlings of WT, *snrk1a*, and *snrk1bc* grown under normal condition (e) or subjected to starvation (f). **(g–l)** Representative plants of WT, *snrk1a*, and *snrk1bc* at maturity in the greenhouse. Aboveground shoot (g), roots (h), shoot and root biomass (i-j), reproductive or yield traits, number of seeds per panicle (k) and weight of 100 seeds (l). **(m-p)** Disease response of detached leaves following inoculation with *M. oryzae* strain Guy11. Two types of assays using spotting (m) or spray (n) of the inoculum is shown in which YT16 serves as the susceptible check. Lesion area in the mutants in spotting (o) and spray (p) assays is compared to WT Kitaake. Significant differences (*p*<0.05) by one-way ANOVA followed by Tukey’s HSD test are shown by small betters. Error bars represent standard deviation (SD). n= 10 (a-f), 20 (g-l), 18 (m, o), and 27 (n, p).

SnRK1 plays an important role in the defense against pathogens (Filipe et al., 2018). Therefore, we analyzed the disease response in *snrk1* mutants by spotting or spraying detached leaves with rice blast fungus, *Magnaporthe oryzae* strain Guy11, that causes diamond-shaped lesions with gray center (sporulating lesions) leading to leaf blast disease in susceptible cultivars such as YT16. In both assays, *snrk1* mutants showed enhanced susceptibility compared to WT Kitaake (**Fig. 1m-p**). As expected, YT16 showed diamond-shaped, coalescing lesions with surrounding chlorosis in the spotted area. The WT (cv. Kitaake), on the other hand, showed small, round lesions confined to the spotted area, a characteristic of hypersensitive response consisting of localized cell death. The *snrk1* mutants, however, showed larger, diamond-shaped lesions, consisting of clear gray centers and a significant increase in the lesion size (**Fig. 1m,o**). Similar results were obtained in the spray assay with significant differences in lesion areas between WT and *snrk1* mutants (**Fig. 1n,p**).

In summary, growth defects in *snrk1* mutants in the energy-sufficient and energy-deficient states and their enhanced susceptibility to leaf blast disease indicate regulatory roles of SnRK1 signaling in normal growth processes, metabolic stress, and disease response in rice. To validate the effect of *snrk1* mutation on plant phenotype, two additional mutants representing the second allele of *ossnrk1*α*a* and *ossnrk1*α*bc* each were analyzed. These mutants referred to as *snrk1a.2* and *snrk1bc.2* showed similar phenotypic defects in the seedlings cultivated in normal or starvation conditions on MS media and mature plants grown in the greenhouse as well as enhanced susceptibility to *M. oryzae* (**Fig. S4**).

### SnRK1 plays a regulatory role in energy-sufficient state

As described above, *snrk1* mutants displayed growth defects under normal and starvation conditions, suggesting a broad role of SnRK1 signaling in rice seedling growth. To better understand the underlying processes, we conducted RNA-seq analysis on WT, *snrk1a,* and *snrk1bc* seedlings under normal and starvation conditions (**Fig. 2a**). Principal component analysis revealed a greater variation of gene expression between *snrk1bc* and WT than between *snrk1a* and WT under normal condition (**Fig. S5a-b**). Accordingly, in comparison to WT, 114 and 1982 differentially expressed genes (p-adj.<0.05) were observed in *snrk1a* and *snrk1bc,* respectively (**Fig. 2b-c; Table S1-S2**), 60 of which were common to the two mutants (**Fig. 2d**). Of these 60 genes, 56 are similarly regulated: 55 are upregulated and 1 is downregulated in both mutants, indicating overlapping roles of *OsSnRK1*α *A, B, C* in SnRK1 signaling.

**Figure 2:**
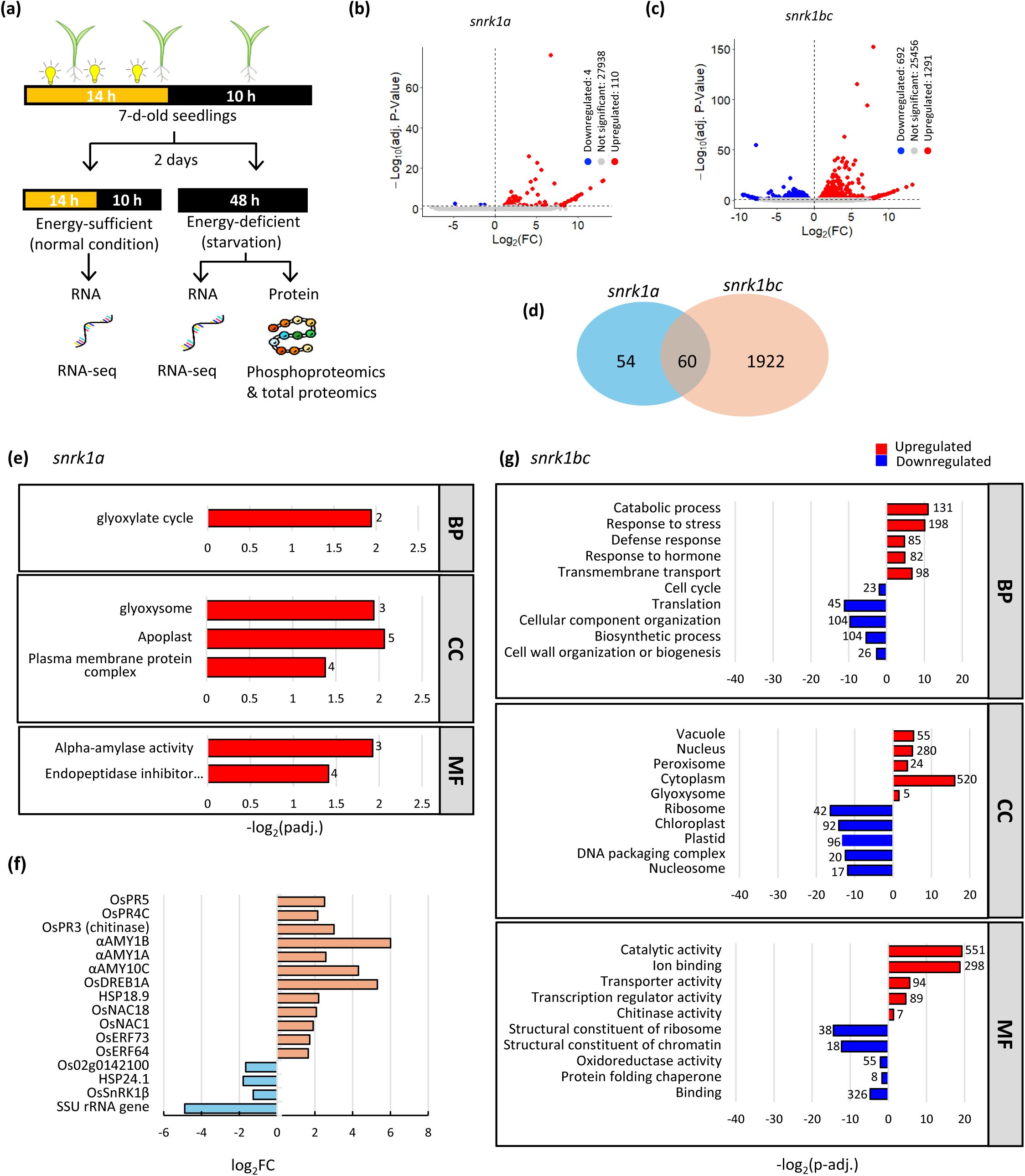
Experimental set up and transcriptomic response of *snrk1* mutants under energy-sufficient (normal) conditions. **(a)** Experimental setup for transcriptomic, proteomic and phosphoproteomic analyses of *snrk1* mutants. 7-d-old seedlings grown in 14 h photoperiod on half-strength MS media were divided into two treatment groups: (i) Energy-sufficient (normal) condition, where seedlings were maintained under 14 h photoperiod for two additional days, and (ii) energy-deficient (starvation) condition, where seedlings were transferred to complete darkness for 2 days (48 hours) starting 12 noon. Total RNA was extracted from seedlings in normal and starvation conditions for transcriptomic analysis and total protein was extracted from seedlings subjected to starvation for proteomic and phosphoproteomic analysis. **(b-c)** Volcano plots showing differentially expressed genes (DEGs) in *snrk1a* (b) and *snrk1bc* (c) compared to WT in energy-sufficient condition. Red and blue dots indicate upregulated and downregulated genes, respectively (p-adj.≤0.05). **(d)** Venn diagram showing unique and overlapping DEGs between *snrk1a* and *snrk1bc* in 9-d-old seedlings under energy-sufficient state. **(e)** Enrichment of Gene Ontology (GO) terms in upregulated DEGs in *snrk1a* mutant. Enriched biological process (BP), cellular component (CC), and molecular function (MF) terms are shown. **(f)** Log fold change of selected DEGs (p-adj.≤0.05, |log FC| > 1) in *snrk1a* based on RNA-seq analysis, including upregulated stress-related transcription factors (e.g., *OsERF64*, *OsNAC1*, *OsDREB1A*), pathogenesis-related genes (*OsPR4C*, *OsPR3*), α-amylase genes (α*AMY10C*, α*AMY1A*, α*AMY1B*), and the downregulated genes, including the β-subunit of *OsSnRK1*, *SSU rRNA*, *HSP24.1*, and *Os02g0142100*. **(g)** GO enrichment analysis of DEGs in *snrk1bc* mutant. Red and blue bars represent upregulated and downregulated GO terms, respectively (p-adj.≤0.05).

The gene ontology (GO) enrichment analysis in *snrk1a* showed association of upregulated genes with glyoxylate cycle in biological process (BP), glyoxysome, apoplast and plasma membrane protein complex terms in cellular component (CC), and α-amylase and endopeptidase inhibitor activities in molecular function (MF) (**Fig. 2e; Table S3**). However, other well-known stress-responsive genes such as *OsERF64, OsERF73, OsNAC1, OsNAC18, HSP18.9, OsDREB1A* were also upregulated (p-adj.≤0.05, log_2_FC>1), along with three pathogenesis-related genes, *OsPR3*, *OsPR4C* and *OsPR3* (chitinase) and 3 α*AMY* genes (α*AMY10C,* α*AMY1A, and* α*AMY1B*). Only 4 genes were significantly downregulated (p-adj.≤0.05, log_2_FC<1) in *snrk1a* mutant that included β-subunit of *OsSnRK1,* small subunit (SSU) *rRNA* gene, *HSP24.1*, and *Os02g0142100* (**Fig. 2f**). In *snrk1bc*, on the other hand, several BP terms were associated with upregulated or downregulated genes (**Table S4**). Notably, stress-related BP were upregulated, while growth-related BP were downregulated. The upregulated processes included catabolic process, response to stress, and defense response, and downregulated processes included cell cycle, translation and biosynthetic process. Accordingly, CC terms associated with upregulated genes included vacuole and peroxisome, and those with downregulated genes included ribosome, plastids and nucleosome. Similarly, MF terms, catalytic activity, ion binding, and chitinase activity were associated with upregulated genes and structural constituents of ribosome and chromatin were associated with downregulated genes (**Fig. 2g**). Taken together, these results indicate that under normal condition, SnRK1 is involved in promoting growth-related processes and suppressing stress-related processes. Collectively, the phenotypic and transcriptomic analyses indicate an important role of SnRK1 in normal growth and development. Higher number of deregulated processes in *snrk1bc*, especially the downregulation of growth processes, aligns with the phenotype of a greater reduction of shoot and root biomass in *snrk1bc* mutant compared to *snrk1a* mutant (**Fig. 1a-b**).

### Starvation-response in *snrk1* mutants is deregulated

SNF1/SnRK1 signaling plays a central role in adaptation to stress and starvation in yeast and Arabidopsis (Baena-González & Sheen, 2008; Coccetti et al., 2018; Hedbacker & Carlson, 2008; Polge & Thomas, 2007). To identify starvation-triggered processes in our *snrk1* mutants, we compared the transcriptome of 9-d-old seedlings under extended darkness, representing starvation/energy-deficient state with that in the normal/energy-sufficient state (**Fig. 2a**). Principal component analysis showed significant differences between the transcriptome of *snrk1a*, *snrk1bc*, and WT under starvation (**Fig. S5c-d**). A total of 3827 genes were differentially expressed (p-adj.≤0.05) in WT during starvation, while *snrk1a* and *snrk1bc* mutants showed only 386 and 843 differentially expressed genes, respectively (**Fig. S6; Table S5-S7**). Hierarchical clustering of these genes showed upregulated (2397 genes) and downregulated (1430 genes) clusters in WT, representing starvation-induced and repressed processes. Starvation-induced genes (cluster 1.1, 1.2) are associated with stress and defense response and starvation-repressed genes (cluster 2) with energy-driven growth processes (**Fig. 3a**). In WT, 1048 genes were upregulated ≥4-fold, while only 36 were downregulated ≥4-fold, suggesting starvation response involves marked upregulation of a large set of genes (**Table S5**). In *snrk1* mutants, several genes in each cluster were deregulated as the number of unique genes in each genotype varied and only 47 genes were common, indicating specific roles of *OsSnRK1* kinase subunit genes and a unique set of upregulated and downregulated processes in each genotype (**Fig. 3b**).

**Figure 3:**
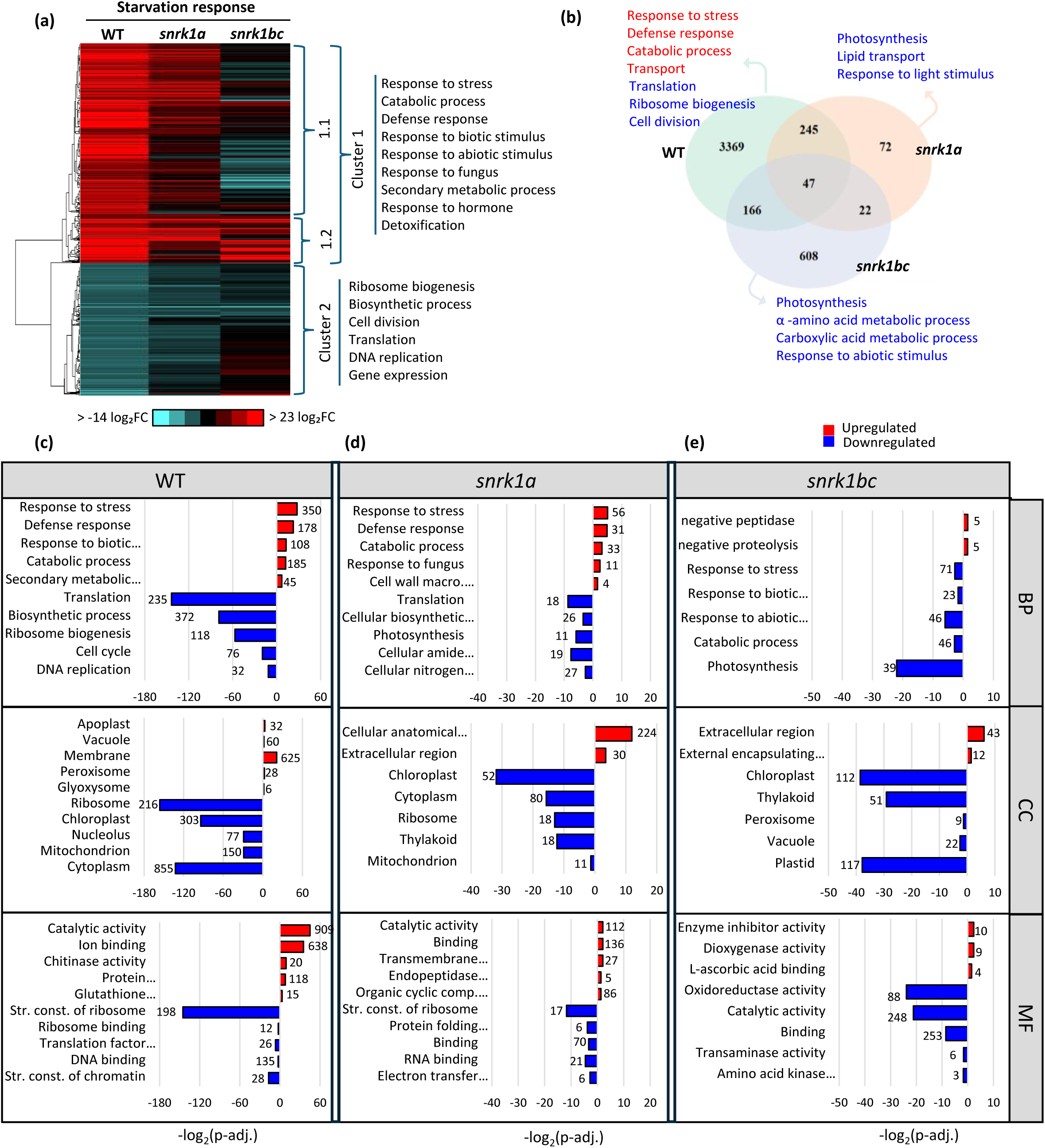
Transcriptomic changes in *snrk1* mutants under starvation. **(a)** Hierarchical clustering of 3827 differentially expressed genes (DEGs) (p-adj.<0.05) in 9-d-old seedlings of WT, *snrk1a*, and *snrk1bc* in starvation. The upregulated and downregulated gene clusters in the WT are shown as cluster 1 and 2, respectively. Cluster 1 is divided into 2 sub-clusters, 1.1 and 1.2, to indicate deregulated genes (1.1) or unaltered genes (1.2) in *snrk1bc* mutant. Major GO terms associated with the clusters are indicated. **(b)** Venn diagram showing the number of unique and common DEGs among WT, *snrk1a*, and *snrk1bc* during starvation. Gene Ontology (GO) processes associated with the unique genes in each genotype are indicated with upregulated GO terms shown in red and downregulated in blue. **(c-e)** Functional enrichment analysis of DEGs showing enriched gene ontology (GO) terms in Biological Process (BP), Cellular Component (CC), and Molecular Function (MF) in WT (c), *snrk1a* mutant (d) and *snrk1bc* mutant (e). Red and blue bars indicate upregulated and downregulated terms, respectively, with numbers of genes associated with each term indicated on each bar (p-adj. ≤0.05).

In the WT, stress and catabolic processes were upregulated and corroborating with the that, CC terms, vacuole, peroxisome, glyoxysome and MF terms, catalytic activity, ion binding, and chitinase activity were associated with upregulated genes during starvation. On the other hand, growth-related processes and CC terms, ribosome and chloroplast, and MF terms, structural components of ribosome were associated with downregulated genes during starvation (**Fig. 3c; Table S8**). The *snrk1a* mutant showed a similar pattern of gene enrichment, but far lower number of genes were associated with each process (**Fig. 3d; Table S9**). However, *snrk1bc* mutant showed an inverse pattern of starvation-triggered processes. First, cluster 1 consisted of two sub-clusters, 1.1 and 1.2 (**Fig. 3a**). While the enrichment of GO processes was similar in the two sub-clusters, 1.1 generally consisted of deregulated genes (SnRK1-dependent), while 1.2 consisted of unaltered genes (SnRK1-independent) in *snrk1bc* mutant. Next, response to stress and catabolic process were downregulated in *snrk1bc* seedlings and negative regulation of peptidase/proteolysis was upregulated. Accordingly, genes associated with CC terms, chloroplast and peroxisome, and MF terms, oxidoreductase activity and catalytic activity were downregulated (**Fig. 3e; Table S10**). These findings corroborate with Arabidopsis studies that showed *snrk1* mutants are unable to turn on stress signaling or respond to starvation (Baena-Gonzalez et al., 2007; Henninger et al., 2022; Pedrotti et al., 2018). These findings also provide mechanistic cues for growth defects in *snrk1* mutants that display abnormally induced starvation-triggered processes in energy-sufficient state, potentially creating a paucity of energy supply, compromising the energy-driven growth processes. It should be noted that *SnRK1* genes are not transcriptionally induced during extended darkness. However, the two rice homologs of *DARK INDUCIBLE 6/ASPARAGINE SYNTHASE 1* (*DIN6/ASN1*), a major marker of Arabidopsis SnRK1 signaling are differentially regulated, and *OsASN1* is induced while *OsASN2* is repressed during starvation (**Fig. S7**).

### Differential phosphorylation of peptides in *snrk1bc* mutant

To identify phosphosites regulated by SnRK1 signaling, phosphoproteomic analysis was carried out on seedlings under starvation. Additionally, total proteomic analysis was carried out to identify the effect of SnRK1 signaling at the proteome level and to correlate protein phosphorylation with protein abundance. We selected *snrk1bc* mutant for this analysis based on the phenotypic and transcriptomic data that showed severe growth defects in its seedlings and a greater number of deregulated genes in comparison to WT. As described above, 9-d-old seedlings exposed to 48 h of continuous darkness were subjected to proteomic and phosphoproteomic analysis (**Fig. 2a**). A comparison of *snrk1bc* vs WT, showed 918 differentially abundant peptides and 248 differentially abundant phosphosites representing 220 proteins (**Fig. 4a-b; Table S11 -S12**). Fifty-two of these proteins were common in the two datasets (**Fig. 4c**), which could be used to understand the effect of phosphorylation on protein abundance. The enrichment analysis of 352 upregulated and 566 downregulated peptides showed upregulation of growth-related processes such as ribosome biogenesis and biosynthetic processes related to cellulose, lipid, amino acids, carbohydrate, and downregulation of energy generation processes such as glucose and ATP metabolic processes, catabolic process as well as response to stress (**Fig. 4d**). Among 248 differentially abundant phosphosites, 102 showed reduced phosphorylation (downregulated) and the remaining 146 showed enhanced phosphorylation (upregulated) in *snrk1bc* mutant (**Fig. 4b; Table S12**). These phosphosites are associated with a functional network consisting of metabolic, growth, and stress processes (**Fig. S8; Table S13**). Notably, upregulated phosphosites are associated with growth processes such as cell division, ribosome biogenesis, and cellulose biosynthesis, and the downregulated phosphosites with energy generation such as glucose metabolic process, ATP generation, and membrane transport (**Fig. 4e; Table S13**). In summary, both proteomic and phosphoproteomic analysis show upregulation of growth-related processes and downregulation of catabolic and stress-related processes in the *snrk1bc* mutant during starvation.

**Figure 4:**
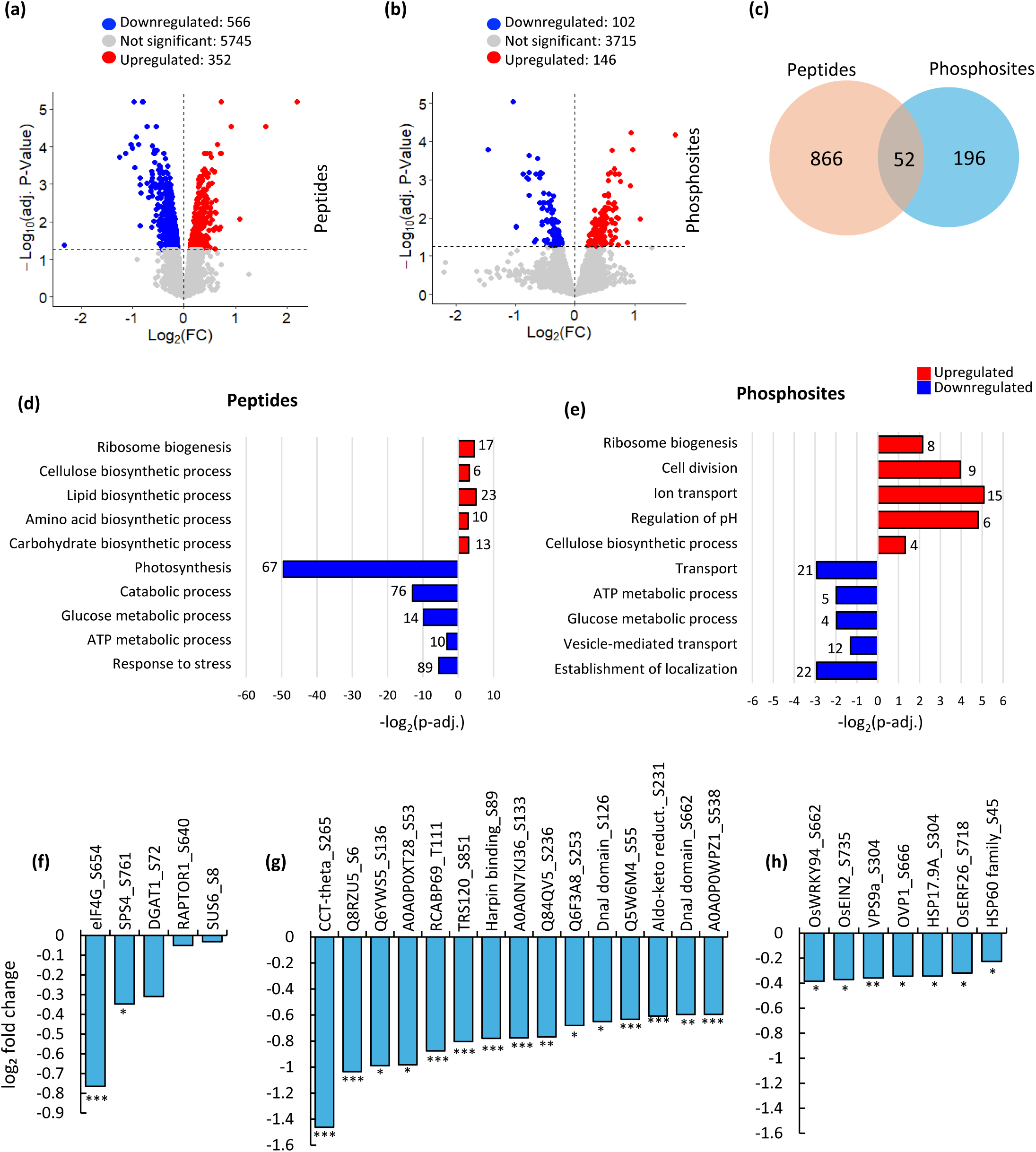
Proteomic and phosphoproteomic analyses of *snrk1bc* mutant under starvation. **(a-b)** Volcano plots of differentially abundant peptides (a) and phophosites (b) in *snrk1bc* compared to WT. Each dot represents a peptide/phosphosite, with the x-axis indicating log fold change and the y-axis showing significance. Differentially abundant peptides/phosphosites (p-adj.<0.05) are shown in red for upregulated, blue for downregulated, and grey for non-significant change. (**c**) Venn diagram showing common or unique proteins detected in proteomic and phosphoporteomic analyses. **(d-e)** Enriched gene ontology (GO) biological processes associated with upregulated (red) or downregulated (blue) peptides (d) and phosphosites (e) in the *snrk1bc* (p-adj.<0.05). **(f-h)** Changes in the phosphorylation levels of the known SnRK1 targets (f), novel targets that show at least 1.5 fold change (g), and selected targets identified among downregulated phosphosites in *snrk1bc* (h). Significance of downregulation for each phosphosite is indicated (*p-adj. ≤ 0.05, *\*\**p-adj. ≤ 0.01, *\*\*\** p-adj. ≤ 0.001).

Of the known SnRK1 targets, Sucrose Phosphate Synthase 4 (SPS4) (Sugden et al., 1999), eukaryotic translation initiation factor 4G (eIF4G) (Cho et al., 2019), Raptor 1 (Gwinn et al., 2008; Nukarinen et al., 2016), Sucrose Synthase 6 (SUS6) (Luo et al., 2020), and Diacylglycerol O-acyltransferase 1 (DGAT1) (Caldo et al., 2018) were downregulated (**Fig. 4f**). These observations indicate deregulation of SnRK1 signaling in *snrk1bc* mutant. Further, 17 phosphosites were reduced ≤1.5-fold, which included eIF4G and putative novel targets: Trafficking Protein Particle Complex II-specific subunit 120 homolog (TRS120), CCT-theta, Harpin binding protein 1, Chlorophyll a-b binding protein (RCABP69), DnaJ domain containing protein, and Aldo-keto reductase 2, and LOC_Os11g43950 protein (Q2R031) (**Fig. 4g**). These phosphosites could be the direct or indirect (through another intermediary protein) targets of phosphorylation by SnRK1. In the protein abundance data (**Table S11**), RCABC69, Aldo-keto reductase 2, and Harpin binding protein 1 were reduced in *snrk1bc* mutant (p-adj.≤0.05), indicating they are stabilized by phosphorylation. Finally, downregulation of the following phosphosites is also noteworthy: transcription factor, OsWRKY94, two major regulators of ethylene signaling, ETHYLENE INSENSITIVE 2 (OsEIN2) and ETHYLENE RESPONSE FACTOR 46 (OsERF46), a major regulator of membrane transport, VPS9a, and two heat shock proteins, HSP17.9A and HSP60 family protein, and vacuolar H^+^ translocating pyrophosphatase (OVP1) (**Fig. 4h**).

### Glucose metabolic process and energy generation is regulated by SnRK1 during starvation

The network of differentially abundant phosphosites in *snrk1bc* mutant showed distinct clusters of downregulated and upregulated processes including glucose metabolic process and energy generation (**Fig. 5a**). Starch biosynthesis and degradation is regulated by photoperiod. During nighttime, starch is degraded to glucose, which is utilized through glycolysis, citric acid cycle, and oxidative phosphorylation to produce energy (ATP). The phosphosites of key enzymes in glycolysis / gluconeogenesis and Calvin cycle within the cluster were differentially abundant in *snrk1bc* mutant. Specifically, Fructose-bisphosphate aldolase 1 (ALDP), chloroplastic; Sedoheptulose 1,7-bisphosphatase (OsSBPase); Glyceraldehyde-3-phosphate dehydrogenase 1 (GAPC1), cytosolic; and phosphoglycerate kinase were significantly downregulated (p-adj.< 0.001). Similarly, the key enzymes in ATP synthesis: ATP synthase subunit alpha, chloroplastic (atpA), and ATP synthase subunit beta, chloroplastic (atpB) were also significantly downregulated (p-adj.<0.0001). The protein abundance of these phosphosites was reduced (p-adj.≤0.05), indicating SnRK1 mediated phosphorylation leads to their stabilization. Together these analyses indicate that ATP generation through glycolytic process during starvation is regulated by SnRK1 signaling in rice seedlings.

**Figure 5:**
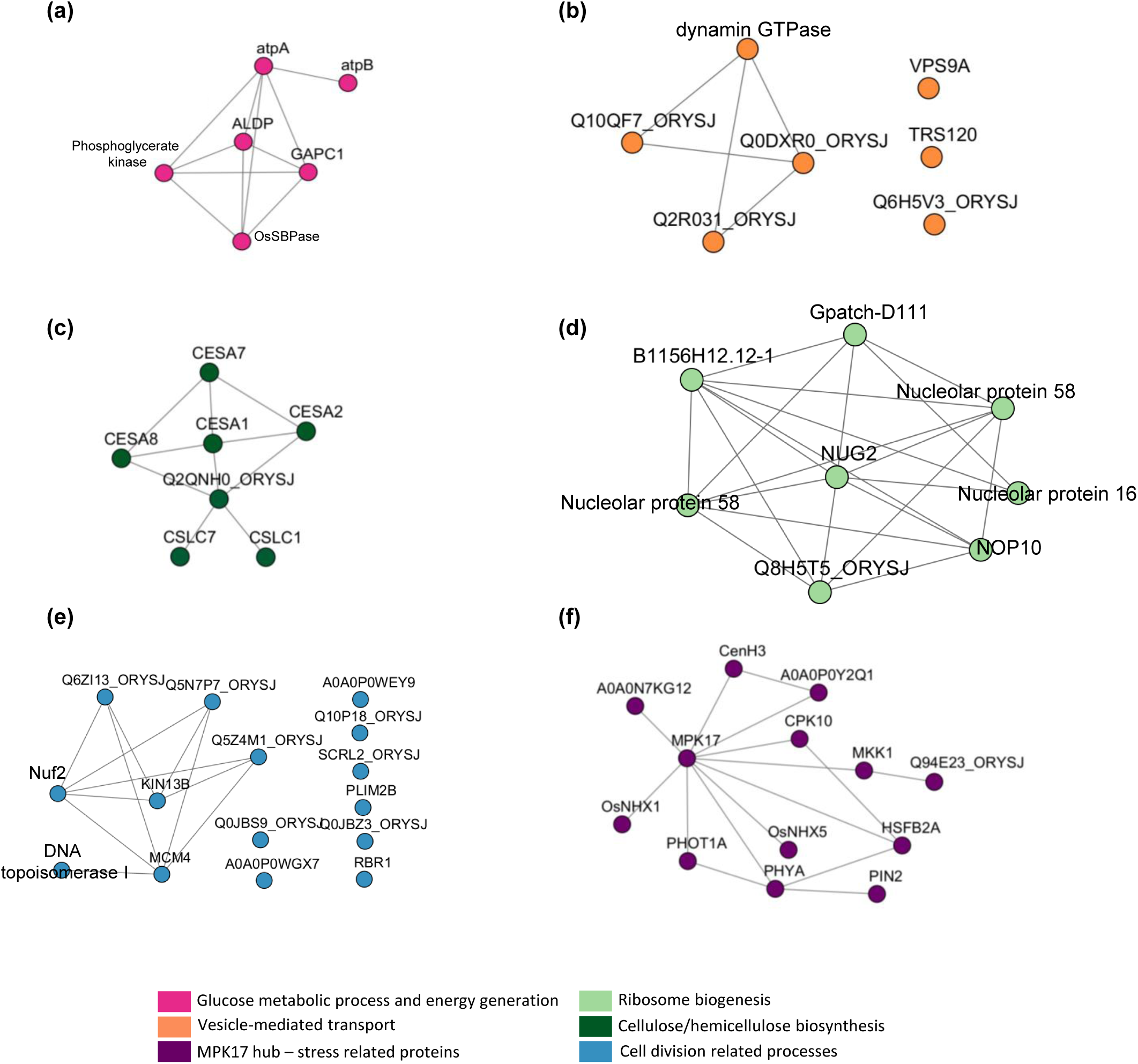
Network of differentially abundant phosphosites in the *snrk1bc* mutant compared to WT under starvation. Each cluster is associated with gene ontology (GO) biological processes color coded in the bottom. **(a-b)** Downregulated phosphosites involved in glucose metabolic process and energy generation (a) or vesicle-mediated transport (b). **(c-e)** Upregulated phosphosites involved in growth processes: cellulose biosynthesis (c), ribosome biogenesis (d), and cell division-related processes (e). **(f)** Upregulated phosphosites associated with stress-related protein network with MPK17 as the hub.

### A potential role of SnRK1 signaling in membrane trafficking

Membrane trafficking is necessary for the integrity of the cell at all points of growth regardless of normal or stress conditions. It involves export and import of material and the movement of cargo to cellular compartments through complex endomembrane networks including vesicles. During energy-deficiency, membrane trafficking is altered to conserve and generate energy from cellular components, inducing autophagy (Feyder et al., 2015; Gill et al., 2021; Zeng et al., 2023). Phosphosites associated with vesicle-mediated transport were affected in *snrk1bc* mutant (p-adj.*<*0.05). Of the 7 downregulated phosphosites in this process (**Fig. 5b**), two are well-known components of vesicle generation and tethering to the target compartment: Vacuolar protein sorting-associated protein 9A (VPS9a) and Trafficking protein particle complex II (TRAPPII)-specific subunit 120 (TRS120) homolog.

VPS9a is a major GDP-GTP exchange factor (GEF) and functional activator for Rab5a small GTPase that regulates initial steps of endocytosis (Carney et al., 2006; Nielsen et al., 2008). For example, in rice endosperm, the transport of dense vesicles from Golgi to protein storage vacuoles (PSV) is regulated by Rab5a homolog Glutelin Precursor 4 (GLUP4) and VPS9a homolog GLUP6. The *glup4* and *glup6* mutants of rice accumulate high amounts of storage proteins in the space between invaginating plasma membrane and the cell wall (Fukuda et al., 2011, 2013). Downregulation of VPS9A/GLUP4 in the phosphoproteome of *snrk1bc* mutant indicates a role for SnRK1 signaling in Rab5a/GLUP4-mediated vesicular transport in rice seedlings. Previously Van Leene et al. (2022) proposed Arabidopsis homolog of VPS9a as the true target of SnRK1. Regulation of VPS9a through phosphorylation is also substantiated by the studies that showed other GEFs such as Rho family and Rab35 GEFs are phosphorylated at tyrosine and/or serine residues (Kulasekaran et al., 2015; Patel and Karginov, 2013). In our dataset, VPS9a/GLUP6 phosphorylation of serine residue was affected (p-adj. = 0.006).

Similarly, phosphorylation of serine residue (S851) in TRS120, a TRAPPII-specific subunit was downregulated in *snrk1bc* mutant during starvation (p-adj.<0.001). TRAPPII is a highly conserved regulator of membrane traffic in both endocytosis and exocytosis pathways (Ravikumar et al., 2017). In Arabidopsis, it is essential for cytokinesis or cell plate formation during cell division. Accordingly, Arabidopsis TRS120 null mutants are seedling lethal due to severe cytokinesis defects (Thellmann et al., 2010). Just as in yeast, Arabidopsis TRAPPII complex functions as a GEF for RabA2 clade of plant Rab GTPases (Kalde et al., 2019). In Arabidopsis, TRS120 is subject to phosphorylation by BIN2, an Arabidopsis SHAGGY-like kinase (Wiese et al., 2024). BIN2 is inhibited by TOR pathway involving phosphorylation by Ribosomal Protein S6 Kinase (S6K) (Xiong et al., 2017). Thus, downregulation of TRS120 phosphosite could be explained by the upregulated TOR signaling in *snrk1bc* mutant leading to inactivation of SHAGGY-like kinase(s).

### SnRK1 suppresses TOR signaling and energy-driven growth processes

The upregulated phosphosites in *snrk1bc* mutant represent indirect targets of SnRK1. These were predominantly associated with growth processes such as ribosome biogenesis, cellulose biosynthesis, and cell division (**Fig. 5c-e**), and possibly related to upregulated TOR signaling in the mutant. Ribosome biogenesis and translation are among the conserved TOR-dependent processes controlled by phosphorylation of S6K that in turn phosphorylates ribosomal protein S6 (RPS6) and eukaryotic translation initiation factor eIF3 and eIF5A (Magnuson et al., 2012; Mahfouz et al., 2006; Schepetilnikov et al., 2013). Earlier Nukarinen et al. (2016) reported enhanced phosphorylation of RPS6 and eIF5A in Arabidopsis *snrk1* mutants. By suppressing TOR signaling through phosphorylation of Raptor 1, SnRK1 antagonistically regulates RPS6, eIF3 subunits, and eIF5A. In *snrk1bc*, phosphosites in RPS6 (Q75LR5 and Q6Z3C2) as well as in eIF5A2 (Q2QQ48) and eIF5B (Q0DFG2) were upregulated (*p*<0.05). eIF3B (Q8S7Q0), eIF3G (Q6K4P1), eIF3M (Q0JFH5), eIF4A-1 (P35683), and eIF6 (Q8GVF5) were also upregulated, although not significantly (**Fig. S9**). These observations serve as molecular evidence for the induced TOR activity in *snrk1bc* mutant.

Among the highly significant upregulated phosphosites (p-adj.<0.02; log_2_FC ≥ 0.585), plasma membrane ATPase, auxin efflux carrier component 2 (OsPIN2), and ammonium transporters (OsAMT1) are noteworthy as they indicate that metabolic activities, nutrient uptake and root development are antagonistically regulated by SnRK1 signaling. The plasma membrane ATPase regulate H^+^ gradient and assist in nutrient uptake (Falhof et al., 2016). In fact, 3 plasma membrane ATPases (Q7XPY2, Q8L6I2, Q8L6I1) and the two OsAMT1s (Q6K9G1, Q7XQ12) that play a major role in ammonium uptake were significantly upregulated in *snrk1bc* mutant (**Table S12**). Next, our data shows that cellulose biosynthesis is antagonistically regulated by SnRK1 signaling, and the upregulation of cell division and cellulose biosynthetic process is another evidence of the unchecked TOR activity in *snrk1bc* mutant during starvation. TOR pathway is integral to cell division, and it likely promotes biosynthesis and deposition of cell wall components such as cellulose and hemicelluloses (Calderan-Rodrigues and Caldana, 2024). The phosphosites in 4 cellulose synthases (CESA1, 2, 7, 8) and 2 hemicellulose biosynthesis enzymes (CSLC1, CSLC7) were upregulated in *snrk1bc* mutant (**Table S12**), suggesting an indirect role of SnRK1 in controlling biosynthesis of the cell wall components (**Fig. 5c**). In addition, growth-related processes such as ribosome biogenesis and cell division were upregulated in the phosphoproteome of the *snrk1bc* mutant (**Fig. 5d-e**), which is known to activate protein synthesis and cell proliferation under nutrient-rich conditions. Next, a cluster of phosphosites associated with stress-related proteins was differentially regulated that consisted of mitogen-activated protein kinase17 (OsMPK17) as the hub protein in the module (**Fig. 5f**). OsMPK17 is a negative regulator of defense response against *Xanthomonas oryzae* pv. *Oryzae* (*Xoo)* and it interacts with MAP kinase kinase1 (MKK1), the positive regulator of defense response against *Xoo* (Yang Z et al., 2024; Zhu et al., 2022). While direct interaction of MPK17 and MKK1 has not been shown, MAP kinase cascade has been implicated in the resistance response against pathogens including *Xoo* (Park et al., 2010). This module also contains light receptors, phytochrome A (PhyA) and phototropin 1 (Phot1A) that are phospho-upregulated. PhyA and phot1A sense far-red and blue light, respectively, and promote photomorphogenesis. Autophosphorylation of PhyA in Ser residues is a key mechanism of attenuating PhyA signaling. In rice, Ser (S599) in the hinge region of PhyA is subject to phosphorylation (Zhou et al., 2018). Phosphorylation of the same Ser (S599) is upregulated in *snrk1bc* mutant (**Table S12**), suggesting an indirect role of SnRK1 in regulating PhyA apoprotein. Overall, proteomics and phosphoproteomics align with transcriptomic analysis of *snrk1bc* as each of these omics show upregulation of growth processes in the *snrk1bc* mutant during starvation and point to the role of SnRK1 signaling in conserving energy by suppressing growth and inducing catabolic processes during stress.

### SnRK1 phosphorylation motifs in rice

To identify possible SnRK1-dependent phosphorylation motifs, we performed motif analysis on the significantly downregulated phosphosites (p-adj.<0.055) in *snrk1bc* mutant using the MoMo tool with the Motif-X algorithm (Cheng et al., 2019). This analysis revealed serine as the predominant residue targeted by SnRK1 (**Fig. 6a-b**). Further, significant enrichment of phosphorylation motifs was seen, particularly those with a P residue at position +1 (P+1) as a classical SP motif, which is characteristic of SnRK1 phosphorylation sites (Cho et al., 2016; Hu et al., 2022; Hu et al., 2024; Nukarinen et al., 2016). Other motifs identified in our dataset include RxxxS (arginine at position P-4) and SxxxL (leucine at position P+4) (**Fig. 6a**), which align with previously reported SnRK1 target motifs characterized by basic residues (R, K, or H) at position P-4 or a hydrophobic residues (M, L, V, I, or F) at position P+4, respectively (Van Leene et al., 2022). We also identified a new motif consisting of an alanine (A) at P-1 (**Fig. 6a**). Overall, this analysis shows that SnRK1 phosphorylation motifs are generally conserved between Arabidopsis and rice and share similarities with AMPK phosphorylation motifs, RxxxS and SxxxL (Hardie et al., 2016).

**Figure 6:**
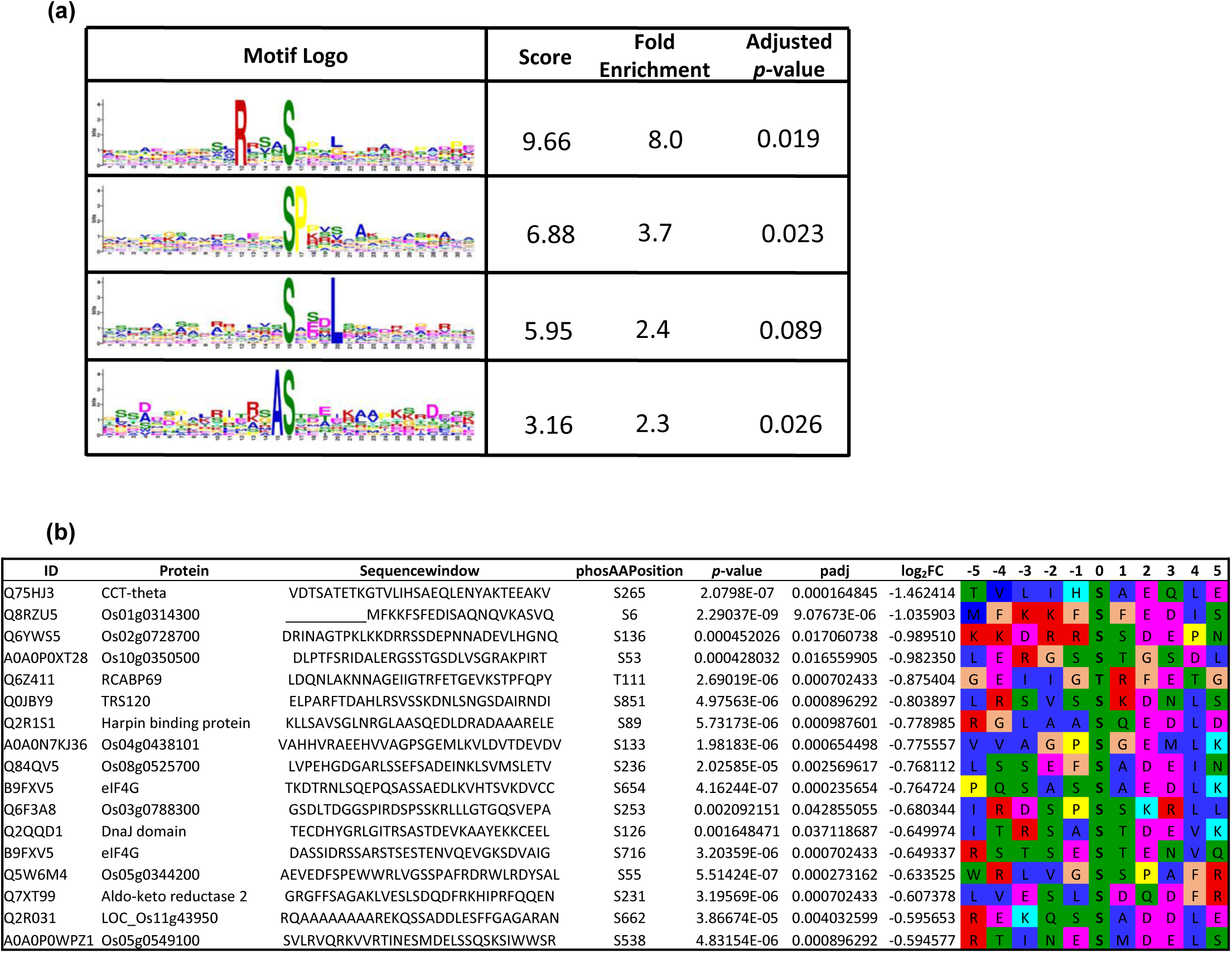
Phosphorylation motif of SnRK1 in rice. **(a)** Motif enrichment in significantly downregulated phosphosites (p-adj.<0.055) in the *snrk1bc* mutant using the MoMo tool with the Motif-X algorithm (*p*-value ≤0.01, at least 10 occurrences). **(b)** sequence motifs in the selected phosphopeptides (p-adj.<0.055; log FC<0.585) in the *snrk1bc* mutant.

## DISCUSSION

Our integrative phenotypic and omics analyses indicate a key role of SnRK1 signaling in regulating growth and adaptive responses in rice. Rice *snrk1* mutants accumulated lower biomass in the seedlings and mature plants and produced a lower number of seeds per panicles (**Fig. 1**). Notably, growth retardation was stronger in *snrk1bc* mutant at early vegetative stage represented by 9-d-old seedlings. However, based on the phenotype of plants at maturity, *snrk1a* mutant showed a greater retardation in the overall growth of shoot and root. These results indicate tissue specificity and functional specialization of *OsSnRK1*α genes. Earlier, Lu et al. (2007) reported retarded germination and shorter shoot and root lengths in 10-d-old seedlings of the *ossnrk1a* mutant. However, the role of other *OsSnRK1*α homologs (*OsSnRK1*α*B* and *OsSnRK1*α*C*) and the role of SnRK1 in other developmental stages (*e.g.* adult growth) is not explored well. Concurring with Lu et al., we found slower elongation of shoot and root from the germinating *snrk1* seeds, with a stronger retardation observed in *snrk1bc* mutant (**Fig. S10**). Next, growth retardation and reduced seed set in our *snrk1a* and *snrk1bc* mutants phenocopy rice plants overexpressing *OsSnRK1*α*A* that also showed decreased shoot and root biomass and lower seed set (Filipe et al. 2018). Taken together, these findings suggest that a balanced SnRK1 signaling is critical for proper growth and development, and disturbance in SnRK1 signaling network leads to growth defects.

The transcriptome of 9-d-old seedlings provided a mechanistic basis of growth defects in normal condition as it showed upregulation of catabolic process and stress-response and downregulation of growth processes (**Fig. 2**), mimicking the starvation response. Wang et al. (2021) also found upregulation of starvation-triggered genes during energy-abundant state in rice *snrk1* mutants. Growth processes during energy-sufficiency are generally regulated by TOR signaling, which is antagonistic to SnRK1. Thus, it is not clear how starvation-triggered processes are upregulated, and TOR-dependent processes are repressed in *snrk1* mutant during energy-sufficiency. Nevertheless, these observations underscore the role of SnRK1 in controlling growth and development even in energy-sufficient (normal) growth conditions. Further, we observed a greater transcriptomic disturbance in *snrk1bc* seedlings than in *snrk1a* seedlings (**Fig. 2**), correlating with the greater phenotypic defect in *snrk1bc* seedlings in normal condition. However, upregulation of stress-related processes, including the upregulation of *OsPR3, OsPR4*, and *OsPR5*, does not lead to disease resistance in *snrk1* mutants. In fact, susceptibility to blast fungus is enhanced in *snrk1* mutants. Previous studies have reported that *snrk1* mutants show enhanced susceptibility to pathogens and overexpression of *SnRK1A* leads to resistance (Cao et al., 2024; Filipe et al., 2018; Kim et al., 2015; Seo et al., 2011). Therefore, SnRK1 signaling is critical for disease resistance. Concurring with that, a recent study showed SnRK1 mediated regulation of plant immunity in Arabidopsis through phosphorylation of NPR1 (Chen et al., 2025), a central regulator of salicylic acid induced defense response.

As expected, *snrk1* seedlings showed growth defects in the starvation state. Increased yellowing in the culm was observed in both *snrk1* mutants, while reduced biomass (shoot and root) was noted in *snrk1bc* seedlings under starvation (**Fig. 1; Fig. S2-S4**). Accordingly, total chlorophyll content was significantly lower in both *snrk1* mutants, and severe phenotypic aberrations were observed in *snrk1bc* seedlings (**Fig. 1**). These observations further point to the functional specialization of *OsSnRK1*α genes and indicate a more prominent role of *OsSnRK1*α*B* and *OsSnRK1*α*C* in seedling development and its starvation response. Starvation is a major inducer of SnRK1 signaling (Baena-Gonzalez et al., 2007; Henninger et al., 2022; Nukarinen et al., 2016; Pedrotti et al., 2018; Wang et al., 2021). Accordingly, the transcriptome of *snrk1* seedlings under starvation showed a larger set of deregulated genes than in the normal condition and corroborated with the severe defect in the phenotype of *snrk1bc* seedlings (**Figs. 2, 3**). Interestingly, normal growth response of *snrk1bc* mutant mirrored the starvation-response of WT as stress and catabolic processes were upregulated in *snrk1bc* in normal condition while being downregulated during starvation (**Fig. S11**). Starvation generally triggers catabolic and stress-related processes as observed in the WT (**Fig. 3**). Deregulation of these processes in *snrk1bc* suggests that the starvation response in rice seedlings is mostly controlled by *OsSnRK1*α*B* and *OsSnRK1*α*C*.

Proteomics and phosphoproteomics provided further validation of the deregulation of SnRK1 signaling in *snrk1bc* mutant by showing upregulation of peptides and phosphopeptide associated with ribosome biogenesis (**Fig. 4-5**). The upregulation of this TOR-dependent process aligns with the current understanding that SnRK1 is required to check TOR activity during starvation (Baena-González & Hanson, 2017; Dobrenel et al., 2016). The downregulated phosphopeptides in *snrk1bc* represent phosphorylation targets of SnRK1. These include core enzymes in energy generation process, aligning with the consensus that SnRK1 is the hub of metabolic reprogramming in response to starvation (Van Leene et al., 2022; Hu et al., 2022). The presence of two proteins functioning in vesicle-mediated transport (VSP9a and TRS120) among downregulated phosphopeptides implicates SnRK1 in intracellular trafficking and corroborates with the study in Arabidopsis that also found a relation of SnRK1 with multiple proteins functioning in protein trafficking, including AtVPS9a (Van Leene et al., 2022). Our findings together with that of Van Leene et al. align with the emerging evidence in mammalians that implicates SnRK1/AMPK in controlling endocytic membrane traffic and ER stress (Chauhan et al., 2020; Rahmani et al., 2019). We also found downregulation of phosphosites of the key proteins in ethylene signaling: OsEIN2 (Q0D8I9) and OsERF46 (Q6H5V3) (**Table S12**), which are positive regulators of ethylene signaling and play a central role in development and stress response (Muller et al., 2015; Jun et al., 2004). Next, three heat shock proteins (Q2QU06, Q2R031, Q84Q77) and Chaperonin Containing TCP1 Subunit 8 (Q75HJ3; CCT-theta) were downregulated, indicating a role for SnRK1 in regulating chaperon protein functions. Further, phosphorylation of eIF4G (B9FXV5) was downregulated in *snrk1bc* mutant. eIF4G is essential for rice tungro virus to manifest disease in susceptible rice cultivars and a mutation in this gene leads to virus resistance (Lee et al. 2010; Macovei et al., 2018). Finally, OsWRKY94 (Q2QMN4) and the proton pump, OVP1 Q67WN5) were downregulated in the phosphoproteome of *snrk1bc* mutant. These proteins play crucial roles in cold and salinity tolerance (Chen et al., 2018; Zhang et al., 2011). In summary, several stress-related phosphopeptides, most of which also play important roles in developmental processes, are downregulated in *snrk1bc* mutant, indicating a pivotal role of SnRK1 in controlling stress and developmental processes.

We propose the true targets of SnRK1 by searching the literature for SnRK1 targets and checking the phosphorylation motif of the downregulated phosphosites. Earlier, Hu et al. (2022) carried out phosphoproteomics using *snrk1a* mutant to study SnRK1-mediated transport of carbohydrates from sheath to panicles and predicted several SnRK1 targets in rice based on parallel reaction monitoring approach of mass spectrometry. We found 9 phosphosites representing 7 proteins in our dataset that were predicted by Hu et al. to be SnRK1 targets (**Fig. S12**). One of them, Os04g0679400 protein, contained the classical SnRK1 target motif (SP), raising the confidence of prediction. VPS9a showed two phosphosites, one of which contained RxxS motif described as SnRK1 target in Arabidopsis and rice (Van Leene et al., 2022; Nukarinen et al., 2016; Hu et al., 2022), and the other contained VxxxxSxxxL that matches the SnRK1 target motif in Arabidopsis, specifically presence of a hydrophobic residue at P-5 and P+4 (Van Leene et al., 2022). Five other proteins also contained motifs described as putative SnRK1 targets in the literature that included RxxxS, SD, a hydrophobic residue (M, L V or I) at P-5 or P+4, and a basic residue (R) at P-3/-4 (**Fig. S12**).

In conclusion, this study demonstrates that SnRK1 signaling is essential for integrating energy status in growth and stress responses in rice. Loss of SnRK1 function disrupts transcriptional and post-translational regulation of metabolic and defense pathways, leading to enhanced disease susceptibility and growth defects. Our findings lead to a potentially novel role for SnRK1 in regulating membrane trafficking during starvation, as indicated by altered phosphorylation of key vesicle-mediated transport proteins. In addition, novel candidates of SnRK1 were identified through phosphoproteomic analysis; however, their direct phosphorylation by SnRK1 remains to be experimentally validated. Overall, our findings show that SnRK1 signaling is critical for plant adaptation and development under both energy-sufficient and energy-deficient conditions. Further, the transcriptomics, proteomics and phosphoproteomics provides a rich resource for further studies focusing on understanding the SnRK1 signaling network in rice.

## ACKNOWLEDGMENTS

This work was supported by the seed grant from ABI-Arkansas Division of Agriculture, and the voucher program from IDEA National Resource for Quantitative Proteomics, UAMS, Little Rock, Arkansas. We are grateful to Martin Egan and his group for their help with leaf blast assays, Jeffery A. Lewis for help with omics data analysis, and Samuel G. Mackintosh for proteomic and phosphoproteomic analysis.

## AUTHOR CONTRIBUTIONS

VS and MJK designed project. MFC, MJK, and VS analyzed the data. CM prepared samples for RNA-seq and proteomics and carried out initial omics analysis. MFC and VS wrote the paper.

## EXPERIMENTAL PROCEDURES

### Plant material and growth conditions

*snrk1a* and *snrk1bc* mutants were developed by CRISPR/Cas9 targeting of *OsSnRK1α* genes as described earlier (Pathak et al., 2022). For seedling analysis, the homozygous lines of the single mutant of *OsSnRK1αA* (*snrk1a*) and double-mutant of *OsSnRK1αB* and *OsSnRK1αC* (*snrk1bc*) along with WT were sterilized and plated on half-strength MS media (MS½) with 2% sucrose solidified with phytagel (2 g/L). Three – four days after plating, at the S3 stage (when prophyll emerges out of the coleoptile), the seedlings were transferred to glass tubes containing MS½ media without sucrose, solidified with phytagel (1.5 g/L). The seedlings were grown in the growth chamber at 26 °C with optimal light (∼30 PAR). The light was provided from 6 am to 7 pm for a 14 h photoperiod. When the seedlings were at V1 stage (formation of first leaf with collar, ∼7 days post S3 stage), the seedlings were divided into 2 groups: half of them were transferred to complete darkness, mimicking starvation, and the other half were kept under light for 2 additional days, totaling 9 days of growth in 14 h photoperiod, representing normal growth. For the phenotyping of mature plants, 7-d-old seedlings at V1-V2 stage were transferred to the greenhouse in pots (7x7x13 cm) filled with commercial potting mix (Promix LP15, Premier Tech Horticulture) consisting of sphagnum peat moss and perlite (9:1). The plants were grown in randomized block design in the greenhouse and fertilized with iron chelate and Osmocote fertilizer (15N-9P-12K) and treated with insecticide (abamectin) as needed.

### RNA-seq analysis

Total RNA from seedlings was isolated using Trizol reagent (Invitrogen Inc.), treated with DNase I, and quantified by Nanodrop 2000 (Thermo Fisher Scientific, USA). Approximately 3 µg per sample (A260/A280 > 1.9) was sent to Novogene Inc. for 150 bp paired-end directional mRNA (poly A enriched) sequencing, generating 7 – 9 GB raw data per sample on Illumina® HiSeq. Two biological replicates of each genotype/treatment consisting of 2 – 5 seedlings each were used. Clean sequences were mapped against *Oryza sativa* japonica Kitaake genome and differentially expressed genes (DEGs) identified using DESeq2 using a threshold of adjusted *p*-value (p-adj.) ≤ 0.05.

### Chlorophyll content

Based on the method described by Pedrotti et al. (2018), 9-d-old seedlings (bulked) were frozen in liquid nitrogen. 100 mg of frozen tissue was ground in 1 mL of methanol homogenized using a Mixer Mill (MM400; Retsch) and incubated at 60 °C for 30 minutes followed by an additional 10 minutes at room temperature (RT). The extract was clarified by centrifugation on a benchtop centrifuge and 1:10 dilution of the supernatant was used for measuring absorbance at 650 and 665 nm in a spectrophotometer. Total chlorophyll (mg chlorophyll/mL extract) was calculated as: total chlorophyll = A_f[650]_ X 0.025 + A_f[665]_ X 0.005.

### Protein extraction

Snap-frozen seedlings (bulked) were ground in liquid nitrogen, and 0.5 g of the powdered tissue was transferred into 2.0 mL microtubes. The ground tissue was suspended in 0.8 mL of extraction buffer [30% sucrose, 2% SDS, 0.1 M Tris-HCl (pH 8.0), and 5% 2-mercaptoethanol] supplemented with phosphatase inhibitor cocktail [PhosSTOP^TM^ tablet (Roche) and 20 µL of phenylmethylsulfonyl fluoride]. An equal volume of phenol buffered with Tris-HCl (pH 8.8) was added to the mixture, which was then vigorously shaken for 30 seconds. The samples were centrifuged at 15,000xg for 20 minutes at 4 °C, and the upper phenol phase was transferred to fresh microtubes. The remaining tissue was re-extracted with an additional 0.8 mL of buffered phenol and 0.8 mL of extraction buffer, and the phenol phase was combined with the first extraction. The pooled phenol phase was used to precipitate protein with 4 volumes of cold methanol containing 100 mM ammonium acetate, thoroughly mixed and placed in -20°C for 30 minutes. The samples were centrifuged at 15,000xg for 20 minutes at 4 °C, and the supernatant was carefully discarded, and protein pellet was washed with 1 mL of cold 80% acetone, vortexed, and centrifuged at 15,000xg for 10 minutes at 4°C. This washing was repeated twice, and the washed protein pellet was used for proteomic and phosphoproteomic analysis.

### LC-MS/MS for discovery proteomics and phosphoproteomics

Protein samples were processed at the UAMS Proteomics Core Facility (https://idearesourceproteomics.org/). Total protein from each sample was reduced, alkylated, and purified by chloroform/methanol extraction prior to digestion with sequencing grade modified trypsin/LysC. Peptides were labeled with TMT 10-plex isobaric label reagent set (Thermo Scientific), and phosphopeptides were enriched using TiO and Fe-NTA kits. Enriched and unenriched labeled peptides were fractionated by high-pH reversed-phase chromatography and further separated on a XSelect CSH C18 2.5 um resin (Waters) using an UltiMate 3000 RSLCnano system (Thermo Scientific). MS analysis was performed on an Orbitrap Eclipse Tribrid mass spectrometer (Thermo Scientific) using multi-notch MS3 parameters. Proteins were identified and reporter ions quantified by searching the UniprotKB database restricted to *Oryza sativa japonica* using MaxQuant (Max Planck Institute, ver. 2.1.4.0) (Tyanova et al., 2016) with FDR<0.01. Protein IDs were assigned using the Protein Prophet algorithm (Nesvizhskii et al., 2003). TMT reporter ion intensities from unenriched samples were used to assess changes in total protein abundance, while phospho Ser/Thr/Tyr (STY) modifications were identified and quantified from the samples enriched for phosphorylated peptides. Following database search, MS3 reporter ion intensities were normalized using ProteiNorm and variance stabilizing normalization (Graw et al., 2020; Huber et al., 2002). Differential abundance analysis was performed using the limma package with empirical Bayes, smoothing to the standard errors (Ritchie et al., 2015). For phosphopeptides, only sites with localization probability >75% were considered.

### Data analysis

Differentially expressed genes/proteins with adjusted *p*-value (p-adj.) of ≤0.05 were used to select the significant genes and proteins from the data. ggplot and VennDiagram in R were used to generate volcano plots and Venn diagrams. Cluster 3.0 (de Hoon et al., 2004) was used for clustering the differentially expressed genes using hierarchical cluster and Pearson correlation method. Java Treeview (Saldanha, 2004) was used to generate heat maps of the gene clusters. Gene ontology analysis and network analysis was done using String software (Szklarczyk et al., 2019) and the output data was visualized using Cytoscape 3.10.2 (Shannon et al., 2003). Motif analysis was done using the MoMo tool with the Motif-X algorithm (Cheng et al., 2019) using a significance threshold of *p* ≤ 0.01 and a minimum of 10 occurrences per motif.

### Leaf blast disease assays

The *Magnaporthe oryzae* strain Guy11 was grown on 0.8% agar for 10-14 days to allow sufficient sporulation. Once the fungi sporulated, 2 ml of 0.02% gelatin was pipetted onto the plate and a spatula was used to scrape the spores into the solution. The spores were filtered using 1 layer of miracloth and collected in 1.5 ml Eppendorf tubes. The tubes were centrifuged at 5000x g for 5 minutes. The supernatant was removed, and the spores were re-suspended in 1 ml 0.02% gelatin. The spores were diluted and counted using a hemocytometer and diluted to a final concentration of 1 x 10^5^ spores per ml. This diluted suspension of spores was used for disease assay as described below. For leaf spotting, the youngest, fully expanded leaf of 3-4 wk-old plants was cut and placed on 0.8% agar media (adaxial side up). Each leaf was inoculated with 5 drops of 20 µl of spore/gelatin solution and incubated for 5 days. For the leaf spray, 9-d-old seedlings grown in MS½ media in 14 h photoperiod were used. The fully expanded leaf of the seedling was cut and placed on 0.8% agar and inoculated with the spore suspension (described above) using an airbrush sprayer inside a fume hood to ensure uniform coverage. The leaves were placed on the agar plate and incubated for 5 days. For both assays, the infected leaves were scanned using a computer scanner, and the area of each lesion calculated using auto threshold MaxEntropy of the ImageJ program (Schneider et al. 2012). Differences in lesion areas between genotypes were determined by Tukey–Kramer test (HSD) using JMP Statistical Discovery 17 from SAS (Version 13.2.1) using the significance threshold of *p*<0.05.

## SUPPORTING INFORMATION

**File S1:** Protein sequence alignments of the mutant sequences of OsSnRK1A*α*, OsSnRK1B*α*, and OsSnRK1C*α*.

Figure S1: Gene expression analysis of *OsSnRK1*α genes in the *snrk1* mutants.

Figure S2: Phenotypic characterization of *snrk1* mutants under normal and starvation conditions.

Figure S3: Starvation-induced phenotypic defects in *snrk1* mutants: representative images.

Figure S4: Phenotypic characterization of *snrk1a.2* and *snrk1bc.2* (second alleles of *ossnrk1a* and *ossnrk1bc*).

Figure S5: Principal component analysis (PCA) of transcriptomic profiles of *snrk1* mutants and WT under normal or starvation conditions.

Figure S6: Volcano plots showing differential gene expression in WT and *snrk1* mutants under starvation in comparison to normal condition.

Figure S7: Gene expression analysis of Os*SnRK1*α subunit genes and rice homologs of Arabidopsis dark-induced 6 (DIN6).

Figure S8: Protein interaction network associated with differentially abundant phosphosites in *snrk1bc* mutant under starvation

Figure S9: Phosphorylation status of the conserved TOR-S6K targets in *snrk1bc* mutant.

Figure 10: Seed germination on half-strength MS media.

Figure S11: Transcriptomic response of *snrk1bc* under normal condition and WT under starvation.

Figure S12: Conserved SnRK1 target motifs identified in rice.

**Table S1 – S10:** Differentially-expressed genes (DEGs) and the associated enriched gene ontology (GO) terms in 9-d-old seedlings of *snrk1* mutants and WT under normal growth or starvation condition.

**Table S11:** Total proteomic analysis. Differentially abundant peptides in *snrk1bc* mutant compared to WT under starvation.

**Table S12:** Phosphoproteomic analysis. Differentially abundant phosphosites in *snrk1bc* mutant compared to WT under starvation.

**Table S13:** Enriched gene ontology (GO) terms associated with the differentially abundant phosphosites in *snrk1bc* mutant.

