## Supplementary material for "Integrative transcriptomic and phosphoproteomic analysis reveals key components of SnRK1 signaling network in rice": File S1

**Supplementary File S1:** Protein sequence alignments of the kinase subunits of the predicted mutant sequences with the wildtype (WT) of OsSnRK1Aα, OsSnRK1Bα, and OsSnRK1Cα.

Kinase domain, Activation loop, and UBA region are highlighted, ATP binding site is shown in bold, and phosphorylation site in activation loop is highlighted in red

**1. Alignment of the predicted OsSnRK1Aα protein sequence in the *snrk1a* mutant with WT**

**OsSnRK1αA (LOC_Os5g45420/ Os05g0530500)**

```
WT          MEGAGRDGNPLGGYRIGKTLGIGSFGKVKIAEHILTGHKVAIKILNRRKIKSMEMEEKVK      60
Mutant      MEGAGRDGNPLGGYRIGKTLGIGSFGKVKIAEHILTGHKVAIKILNRRKIKSMEMEEKVK      60
            ************************************************************

WT          REIKILRLFMHPHIIRLYEVIDTPADIYVVMEYVKSGELFDYIVEKGRLQEEEARRFFQQ      120
Mutant      REIKILRLFMHPHIIRLYEVIDTPADIYVVMEYVKSGELFDYIVEKGRLQEEEARRFFQQ      120
            ************************************************************

WT          IISGVEYCHRNMVVHRDLKPENLLLDSKCNVKIADFGLSNVMRDGHFLKTSCGSPNYAAP      180
Mutant      IISGVEYCHRNMVVHRDLKPENLLLDSKCNVKIADFGLSNVMRDGHFLKTSCGSPNYAAP      180
            ************************************************************

WT          EVISGKLYAGPEVDVWSCGVILYALLCGTLPFDDENIPNLFKKIKGGIYTLPSHLSPLAR      240
Mutant      EVISGKLYAGPEVDVWSCGVILYALLCGTLHLMTRIFPTFLRK-----------------      223
            ****************************** :  . :*.:::*

WT          DLIPRMLVVDPMKRITIREIREHQWFTVGLPRYLAVPPPDTAQQVKKLDDETLNDVINMG      300
Mutant      ------------------------------------------------------------      223


WT          FDKNQLIESLHKRLQNEATVAYYLLLDNRLRTTSGYLGAEFHESMESSLAQVTPAETPNS      360
Mutant      ------------------------------------------------------------      223


WT          ATDHRQHGHMESPGFGLRHHFAADRKWALGLQSRAHPREIITEVLKALQELNVCWKKIGH      420
Mutant      ------------------------------------------------------------      223


WT          YNMKCRWSPSFPSHESMMHNNHGFGAESAIIETDDSEKSTHTVKFEIQLYKTRDEKYLLD      480
Mutant      ------------------------------------------------------------      223


WT          LQRVSGPQLLFLDLCSAFLTQLRVL       505
Mutant      -------------------------       223
```

**2. Alignments of the predicted OsSnRK1αB and OsSnRK1αC protein sequences in the *snrk1bc* mutant with WT**

**OsSnRK1αB (LOC_Os8g37800/Os8g0484600)**

```
Mutant      MEGNARGGGHSEALKNYNLGRTLGIGSFGKVKIAEHKLTGHRVAIKILNRRQMRNMEMEE      60
WT          MEGNARGGGHSEALKNYNLGRTLGIGSFGKVKIAEHKLTGHRVAIKILNRRQMRNMEMEE      60
```

************************************************************

Mutant KAKREIKILRLFIHPHIIRLYEVIYTPTDIYVVMEYCKFGELFDYIVEKGRLQEDEARRI 120
WT KAKREIKILRLFIHPHIIRLYEVIYTPTDIYVVMEYCKFGELFDYIVEKGRLQEDEARRI 120
************************************************************

Mutant FQQIISGVEYCHSGGSS------------------------------------------- 137
WT FQQIISGVEYCHRNMVVHRDLKPENLLLDSKYNVKLADFGLSNVMHDGHFLKTSCGSPNY 180
************ .

Mutant ------------------------------------------------------------ 137
WT AAPEVISGKLYAGPEVDVWSCGVILYALLCGTLPFDDENIPNLFKKIKGGIYTLPSHLSA 240

Mutant ------------------------------------------------------------ 137
WT LARDLIPRMLVVDPMKRITIREIREHQWFQIRLPRYLAVPPPDTAQQAKMIDEDTLQDVV 300

Mutant ------------------------------------------------------------ 137
WT NLGYEKDHVCESLRNRLQNEATVAYYLLLDNRFRATSGYLGADYQESLERNLNRFASSES 360

Mutant ------------------------------------------------------------ 137
WT ASSNTRHYLPGSSDPHASGLRPHYPVERKWALGLQSRAQPREIMIEVLKALEDLNVCWKK 420

Mutant ------------------------------------------------------------ 137
WT NGQYNMKCRWSVGYPQATDMLDVNHSFVDDSIIMDNGDVNGRLPAVIKFEIQLYKSRDEK 480

Mutant ----------------------------- 137
WT YLLDMQRVTGPQLLFLDFCAAFLTKLRVL* 509

**OsSnRK1αC (LOC_Os3g17980/ Os03g0289100)**

Mutant MLTRTITYCMVSVTHRIHHPSIMNKLFSTAWILRHVLRWFKVKMDGNAKGGGHSEALKNY 60
WT MLTRTITYCMVSVTHRIHHPSIMNKLFSTAWILRHVLRWFKVKMDGNAKGGGHSEALKNY 60
************************************************************

Mutant NLGRT**LGIGSFGKV**KIAEHKLTGHRVAIKILNRRQMRNMEMEEKAKREIKILRLFIHPHI 120
WT NLGRT**LGIGSFGKV**KIAEHKLTGHRVAIKILNRRQMRNMEMEEKAKREIKILRLFIHPHI 120
************************************************************

Mutant IRLYEVIYTPTDIYVVMEYCKFGELFDYIVEKGRLQEDEARRIFQADYIWG--------- 171
WT IRLYEVIYTPTDIYVVMEYCKFGELFDYIVEKGRLQEDEARRIFQQIISGVEYCHRNMVV 180
*********************************************

Mutant ------------------------------------------------------------ 171
WT HRDLKPENLLLDSKYNVKLADFGLSNVMHDGHFLKTSCGSPNYAAPEVISGKLYAGPEVD 240

Mutant ------------------------------------------------------------ 171
WT VWSCGVILYALLCGTLPFDDENIPNLFKKIKGGIYTLPSHLSALARDLIPRMLVVDPMKR 300

Mutant ------------------------------------------------------------ 171
WT ITIREIREHQWFQIRLPRYLAVPPPDTAQQAKMIDEDTLQDVVNLGYGKDHVCESLRNRL 360

Mutant ------------------------------------------------------------ 171
WT QNEATVAYYLLLDNRFRATSGYLGADYQESLERNFNRFASSESASSNTRHYLPGSSDPHA 420

```
Mutant     ------------------------------------------------------------      171
WT        SGLRPHYPVERKWALGLQSRAQPREIMIEVLKALQDLNVSWKKNGQYNMKCRWSVGTQAT      480


Mutant     ------------------------------------------------------------      171
WT        DMLDVNNSFVDDSIIMDNGDVNGRLPAVIKFEIQTRDEKYLLDMQRVTGPQLLFLDFCAD      540


Mutant     ---------      171
WT         FLTKLRVL*      548
```

### Supplementary methods:

**Gene Expression Analysis:** Two micrograms of total RNA were treated with RQ1-RNAse free DNase (Thermofisher Inc.), and one microgram of the DNase-treated RNA was used for cDNA synthesis using PrimeScript RT reagent kit (Takara Bio, CA, USA). The expression analysis was performed using TB green Premix Ex Taq II (Takara Bio, CA, USA) on Bio-Rad CFX 96 C1000 with following conditions: 95°C for 30 sec and 40 cycles of 95°C for 5 sec + 60°C for 30 sec. The product specificity was verified by the melt curve analysis. Rice *UBQ2* gene was used as the internal control. Primer sequences used in this study are given below:

**List of qPCR primers:**

| Gene | Gene ID | Sequence (5' to 3') |
|---|---|---|
| *OsSnRK1A* | Os05g0530500 | Forward: GCAGACTTTGGCTTGAGTAATG<br>Reverse: AACTTCAGGGCCAGCATATAG |
| *OsSnRK1B* | Os08g0484600 | Forward: GCTTGCTGACTTTGGTTTGAG<br>Reverse: CCTCGGGTCCAGCATATAATTT |
| *OsSnRK1C* | Os03g0289100 | Forward: GCAGACTTTGGCTTGAGTAATG<br>Reverse: AACTTCAGGGCCAGCATATAG |
| *OsASN1* | Os03g0291500 | Forward: CGGTTACCTCTACTTCCACTTC<br>Reverse: GTTAGCACGCAGACAGTCATA |
| *OsASN2* | Os06g0265000 | Forward: GGTGGACTGGACTCTTCTTTG<br>Reverse: GCAGCTCTAAGATCAGGAGAAC |
| *OsUBQ2* | Os02g0161900 | Forward: TGGTCAGTAATCAGCCAGTTTG<br>Reverse: CAAATACTTGACGAACAGAGGC |

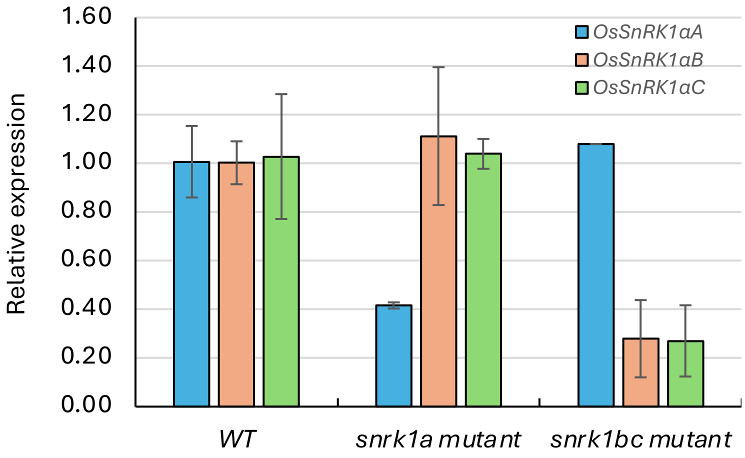


**Figure S1: Gene expression analysis of *OsSnRK1α* genes in rice *snrk1* mutants.** Gene expression of *OsSnRK1A*, *OsSnRK1B*, and *OsSnRK1C* genes determined by real-time qPCR and normalized against *OsUBQ2* gene in 9-d-old seedlings of WT, *snrk1a*, *snrk1bc* mutants grown in 14 h photoperiod on MS½ media. The graph shows expression relative to the value for gene in WT. Mean of 2 – 3 biological replicates are shown with ± standard deviation (SD).

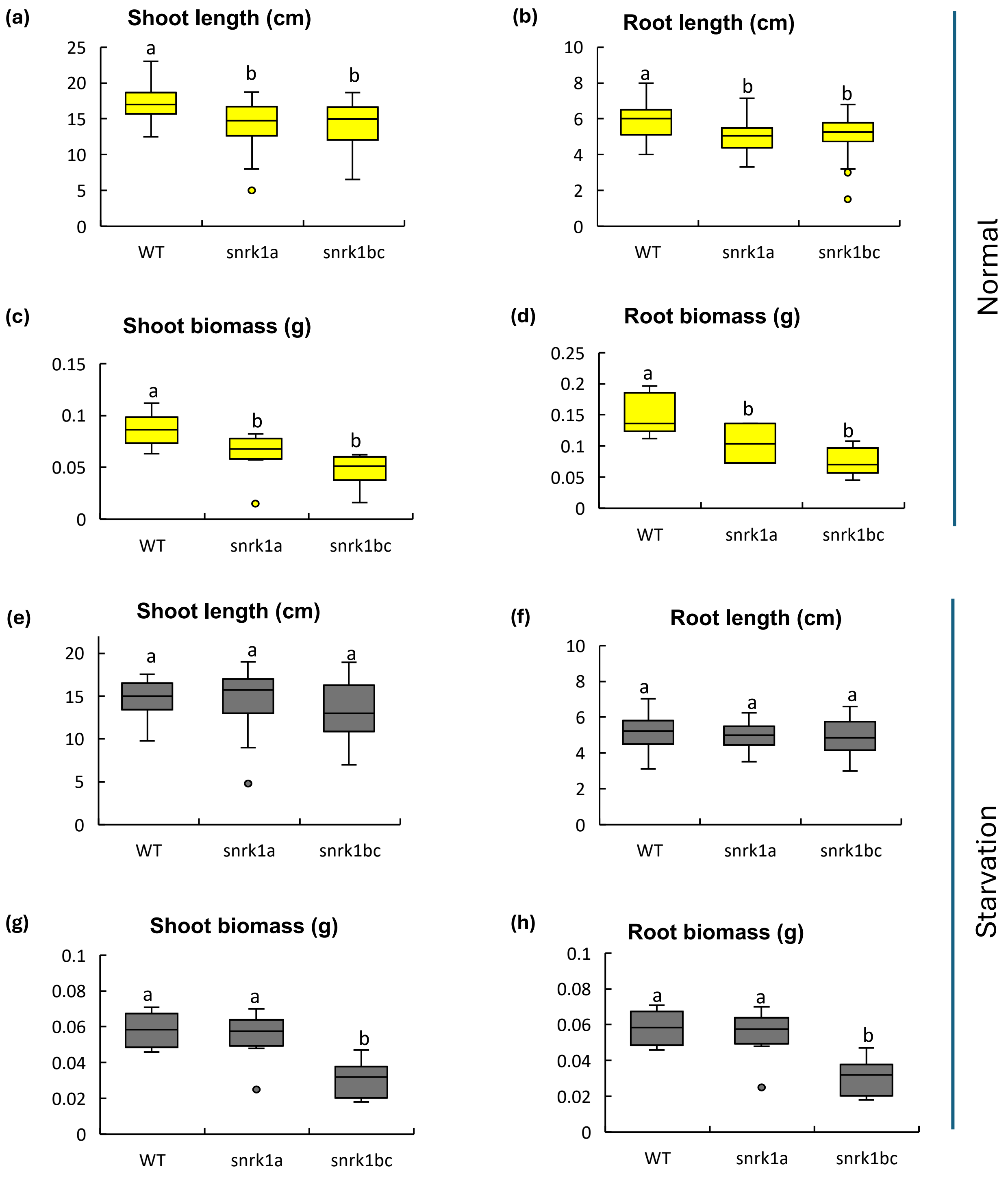


**Figure S2: Phenotypic characterization of *snrk1* mutants under normal and starvation conditions. (a–d)** Boxplots showing shoot length **(a)**, root length **(b)**, shoot biomass **(c)**, and root biomass **(d)** of 9-day-old WT, *snrk1a*, and *snrk1bc* seedlings grown under normal (14-h light/10-h dark) conditions. **(e–f)** Boxplots showing shoot length **(e)**, root length **(f)**, shoot biomass **(g)**, and root biomass **(h)** of seedlings grown for 7 days under normal condition and then exposed to 48 h of continuous darkness (starvation). Different letters indicate significant differences ($p<0.05$) determined by one-way ANOVA followed by Tukey's HSD test on data collected from 10 to 24 seedlings. Error bars represent standard deviation (SD).

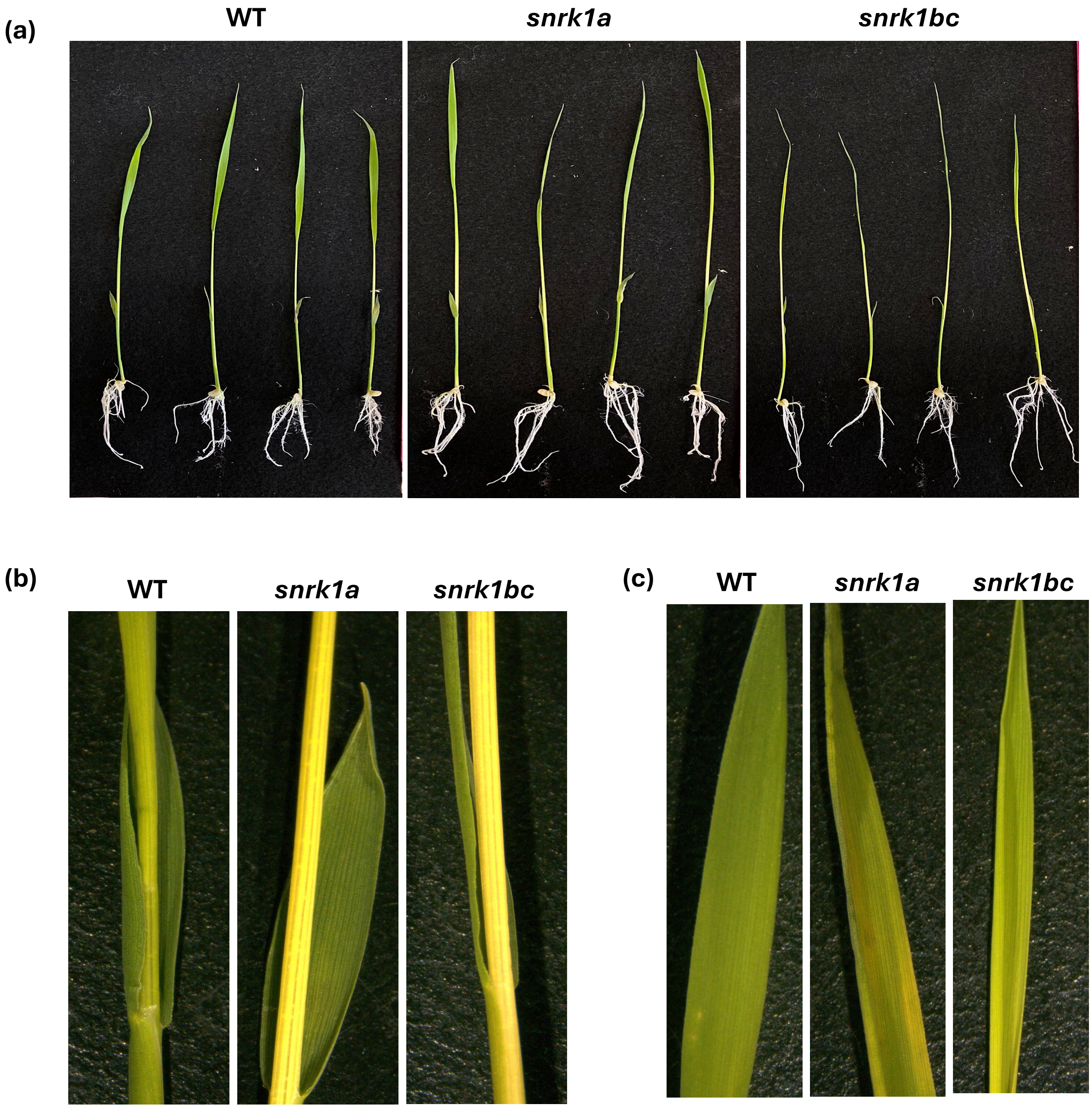


**Figure S3. Starvation-induced phenotype. (a)** Representative images of seedlings of WT, *snrk1a*, and *snrk1bc* after 48 h of continuous darkness (starvation) imposed on 7-d-old seedlings. (**b-c**) close-up of the culm at the first leaf collar (b) and fully expanded leaf (c).

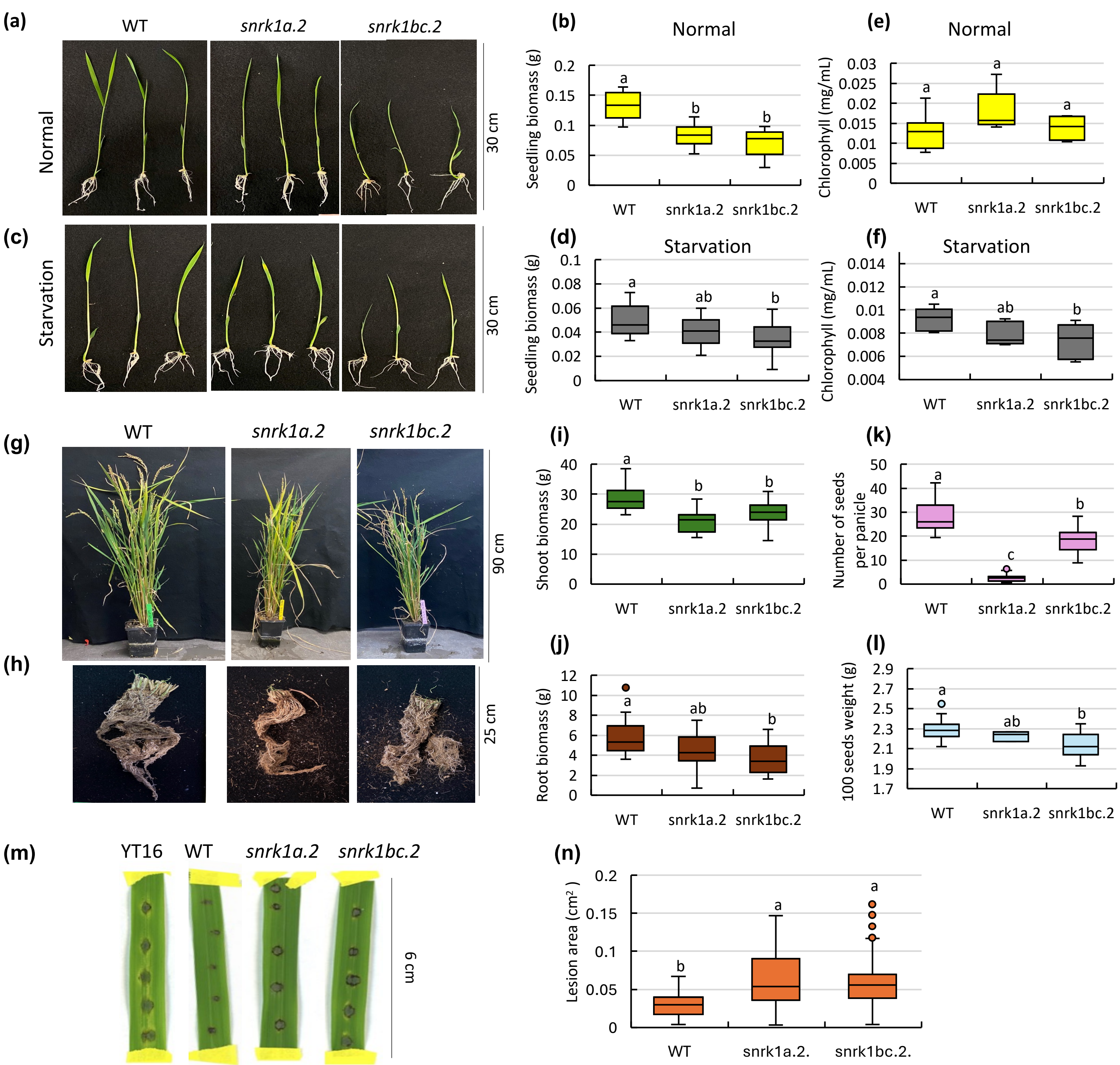


**Figure S4:** Phenotypic characterization of *snrk1a.2* and *snrk1bc.2* (second alleles of *ossnrk1aa* and *ossnrk1abc*). **(a-d)** Seedling phenotype and biomass of 9-d-old seedlings grown on half-strength MS medium under 14-h photoperiod (normal condition) or exposed to 48 h of continuous darkness after 7d of normal growth (starvation). **(e–f)** Chlorophyll content in 9-d-old seedlings under normal or starvation condition. **(g–l)** Plants at maturity in greenhouse conditions. Representative shoot and root images (g-h), shoot and root biomass (i-j), and reproductive traits (k-l). **(m-n)** Disease response of detached leaves following inoculation with *M. oryzae* strain Guy11. YT16 is a susceptible rice. Significant differences ($p$<0.05) by one-way ANOVA followed by Tukey's HSD test is shown by small betters. Error bars represent standard deviation (SD). n= 16 (a-f), n = 20 (g-l), n = 18 (m-n).

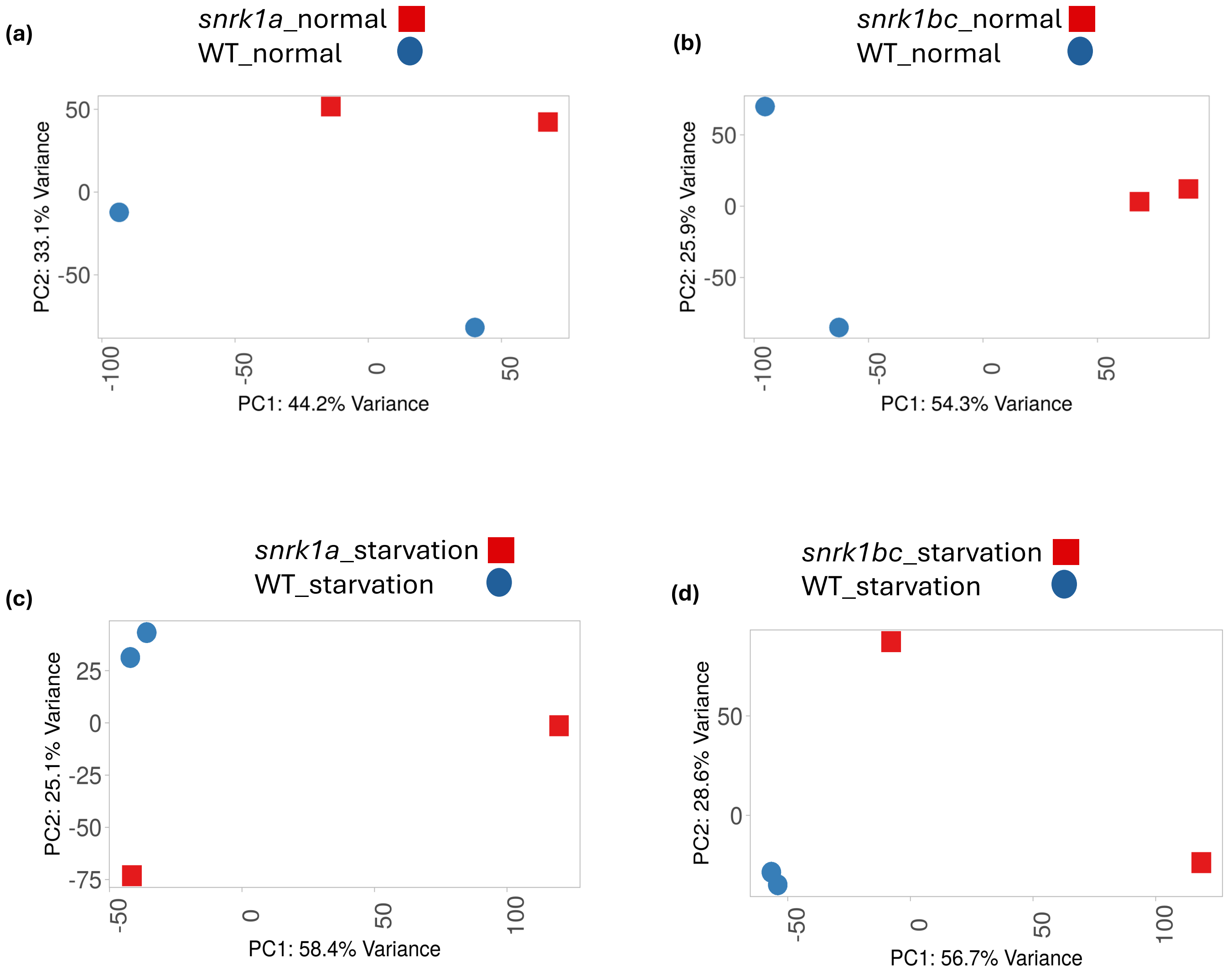


**Figure S5. Principal component analysis (PCA) of transcriptomic profiles of *snrk1* mutants and WT under normal or starvation conditions.** PCA plots of RNA-seq data (normalized gene count) from 9-d-old WT, *snrk1a*, and *snrk1bc* seedlings grown under normal condition (14-h photoperiod) or subjected to 48 h of continuous darkness after 7 days of normal growth (starvation condition). PCA plots were generated on iDEP web-based tool (Ge et al., 2018).

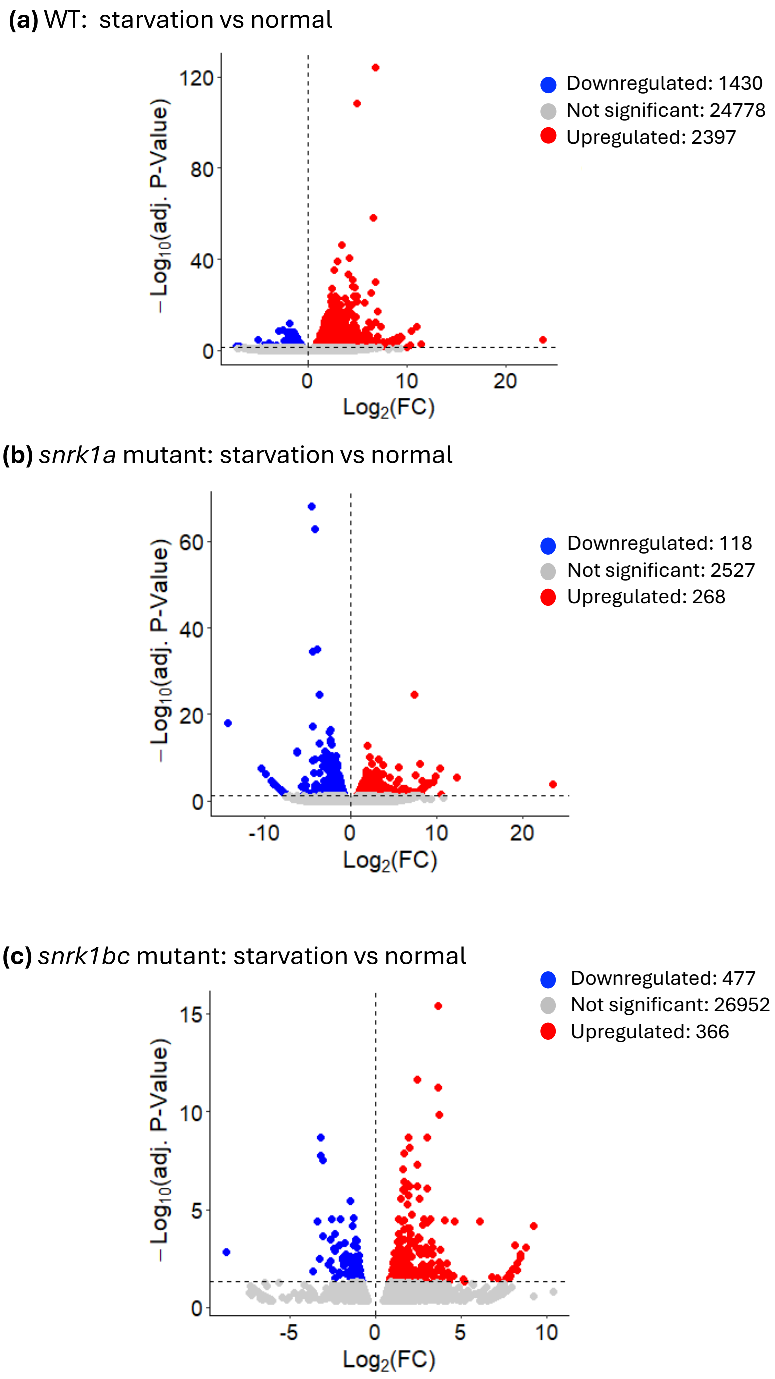


**Figure S6. Differential gene expression in WT and *snrk1* mutants under starvation compared to normal growth.** Volcano plots showing transcriptomic changes in **(a)** WT, **(b)** *snrk1a*, and **(c)** *snrk1bc* seedlings after 48 h of continuous darkness (starvation) versus those grown in normal condition. Differentially expressed genes (DEGs) were identified based on P-adj.< 0.05 and $|log_2FC| > 0$.

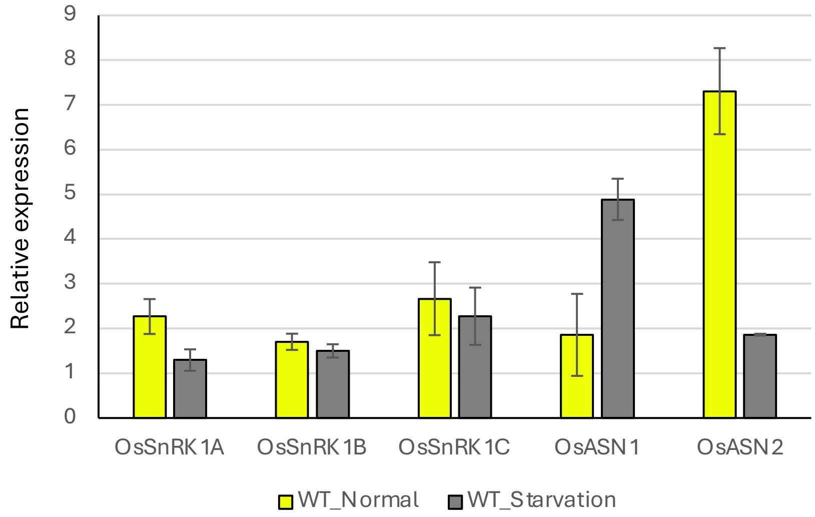


**Figure S7: Gene expression analysis of Os*SnRK1α* subunit genes and rice homologs of Arabidopsis dark-induced 6 (*DIN6*).** qPCR analysis on the rice *OsSnRK1α* paralogs (A, B, C) and the rice homologs of Arabidopsis *DARK INDUCIBLE 6/ASPARAGINE SYNTHASE 1* (*DIN6/ASN1*) (*OsASN1* and *OsASN2*) in 9-d-old seedlings of WT Kitaake seedlings grown in 14 h photoperiod (normal) or exposed to 2 days of continuous darkness after 7 days of normal growth (starvation). Gene expression was normalized against *OsUBQ2* gene (Os02g0161900). Average of 2 – 3 samples (biological replicates) was plotted with standard deviation shown as error bars.

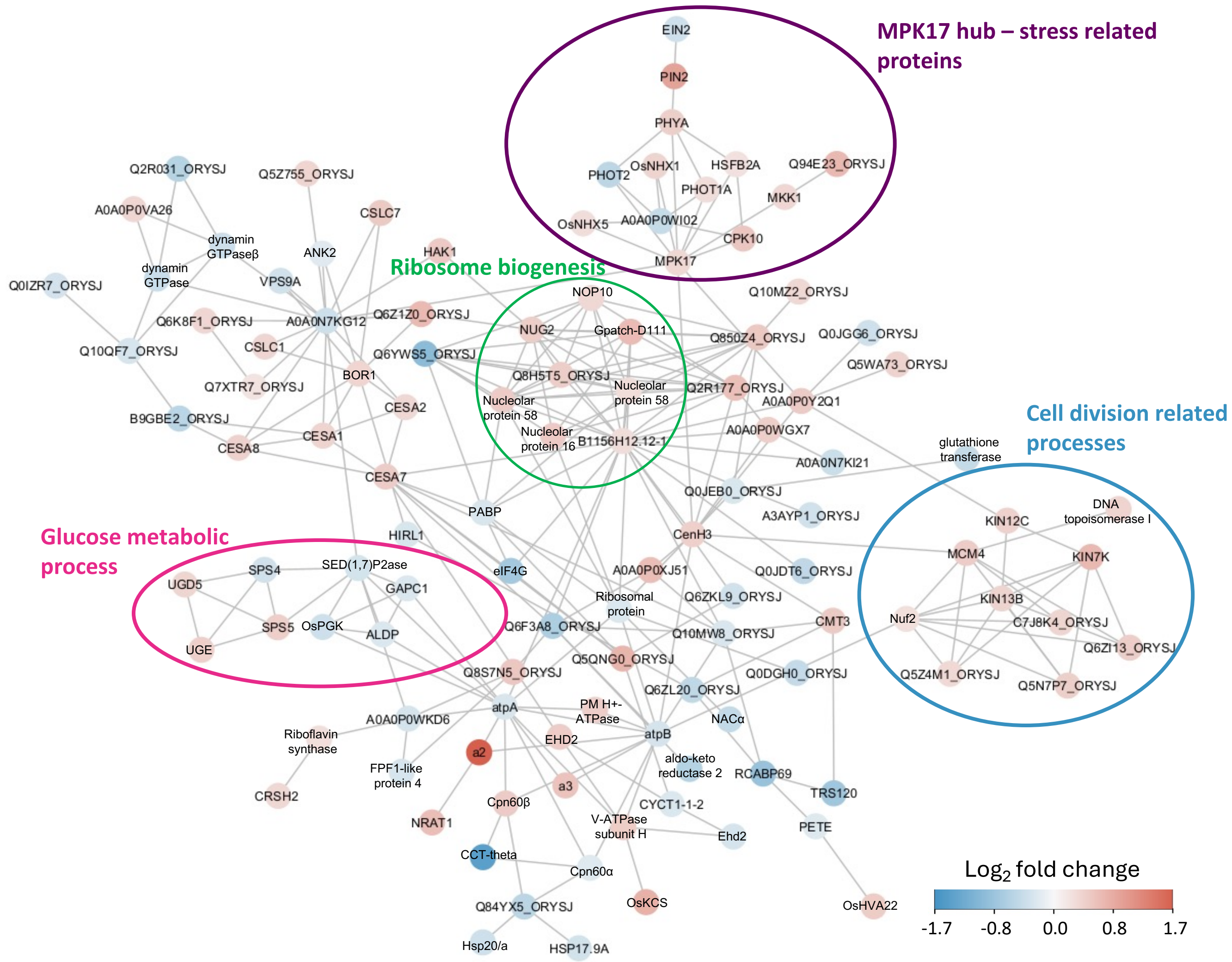


**Figure S8: Protein interaction network associated with differentially-abundant phosphosites in *snrk1bc* mutant under extended darkness.** STRING-based network analysis of significantly abundant phosphosites in *snrk1bc* seedlings under starvation (p-adj.< 0.055; |$\log_2$ fold change| > 0). Nodes represent proteins and edges represent known or predicted protein–protein interactions. Nodes are color-coded by $\log_2$ fold change: red indicates proteins associated with upregulated phosphosites and blue indicates proteins associated with downregulated phosphosites. Selected networks associated with gene ontology (GO) terms are circled.

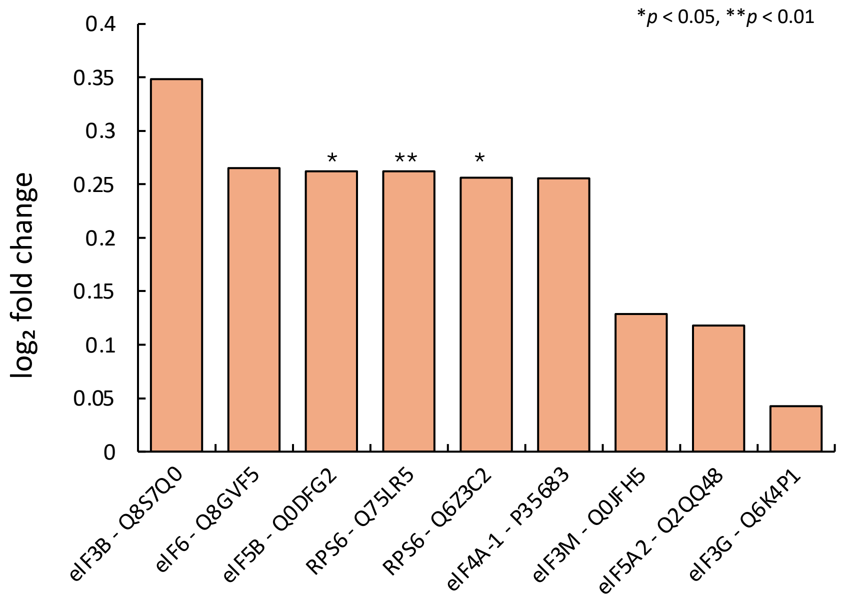


**Figure S9: Phosphorylation status of conserved TOR-S6K targets in *snrk1bc*.** Enhanced phosphorylation ($log_2$ fold change) of ribosomal protein S6 (RPS6) and eukaryotic translation initiation factors (eIFs) in *snrk1bc* compared to WT under starvation. Significance (*p*-value) is indicated as * or **. The rest of the phosphosites show non-significant fold-change.

(a)

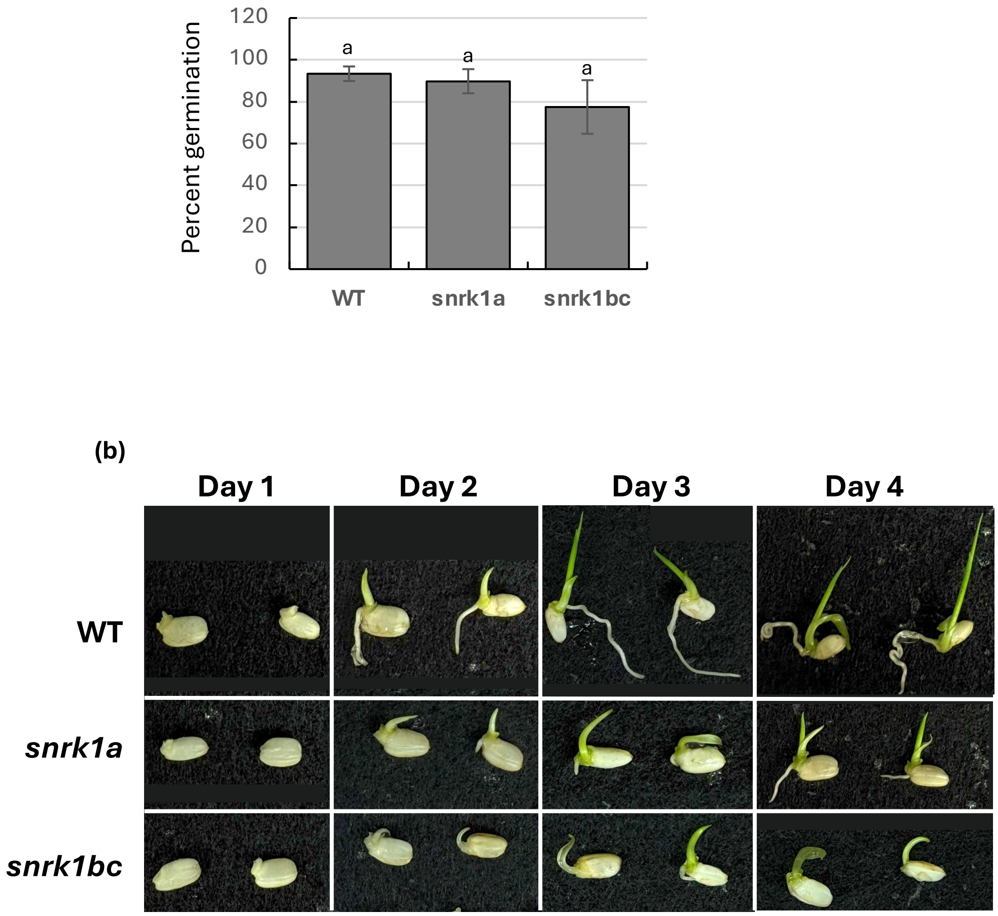


**Figure S10: Seed germination on half-strength MS media. (a)** percent germination based on number of seeds showing emergence of coleoptile after 3 days on the media at 28$^{o}$C in 14 h photoperiod (n=70). **(b)** Representative images of germinating seeds of WT and *snrk1* mutants on the media. Note slower rate of shoot and root elongation in *snrk1* mutants, although percent germination is not significantly different ($p$<0.05) by one-way ANOVA followed by Tukey's HSD test.

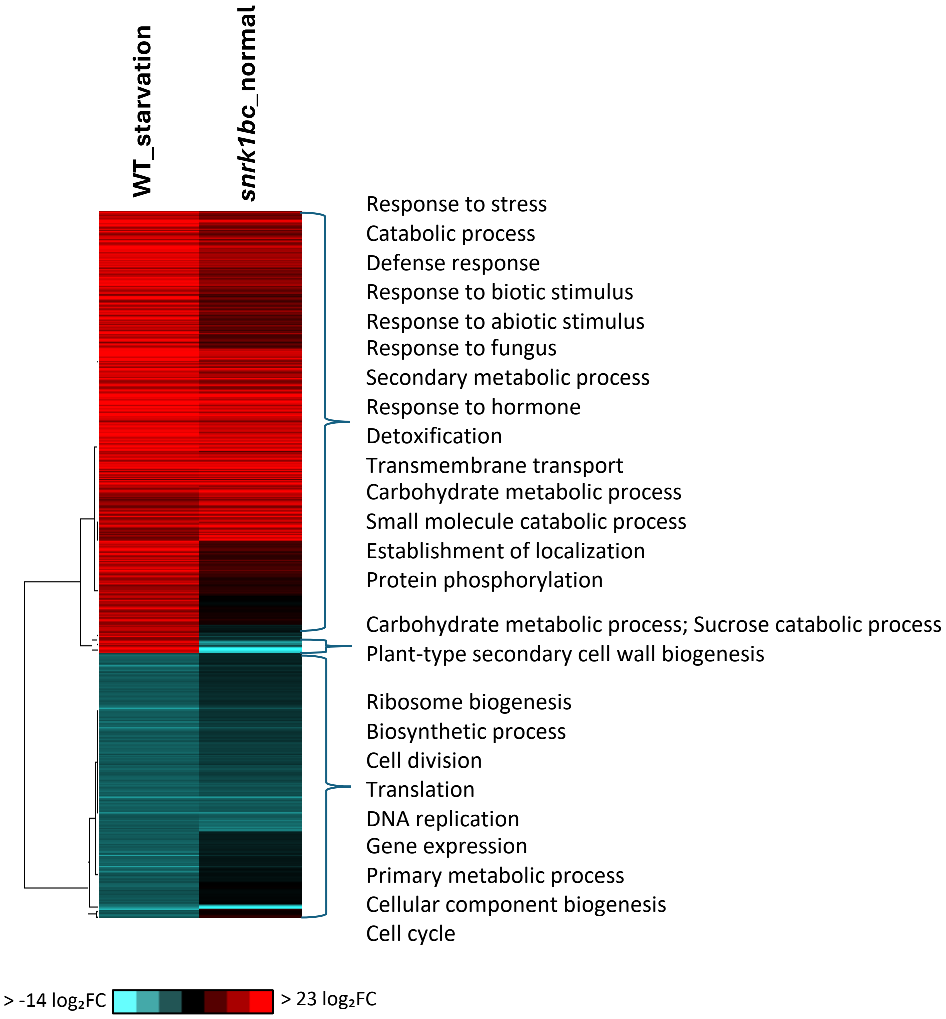


**Figure S11: Transcriptomic profile of *snrk1bc* under normal condition mirrors the transcriptomic profile of WT under starvation.** Hierarchical clustering of differentially expressed genes in WT under starvation (WT_starvation vs WT_normal) and *snrk1bc* mutant under normal condition (*snrk1bc*_normal vs WT_normal). Genes were clustered based on fold-change, with red indicating upregulation and blue indicating downregulation. Associated gene ontology (GO) terms or functional keywords are annotated.

| Proteins | gene name | Sequence window | Phos-AA-Position | *p*-value | p-adj. | $log_2FC$ | -5 | -4 | -3 | -2 | -1 | 0 | 1 | 2 | 3 | 4 | 5 | Reference |
|---|---|---|---|---|---|---|---|---|---|---|---|---|---|---|---|---|---|---|
| Q6Z411 | *RCABP69* | LDQNLAKNNAGEIIG**T**RFETGEVKSTPFQPY | T111 | 0.000003 | 0.000702 | -0.875404 | G | E | I | I | G | **T** | R | F | E | T | G | Hu et al., 2022 |
| Q10NQ3 | *VPS9A* | DVREQKSQTLKASRD**S**DVNLSLKDNFQGPGL | S304 | 0.000092 | 0.006358 | -0.358038 | K | A | S | R | D | **S** | D | V | N | L | S | Hu et al., 2022; Hu et al., 2024 |
| Q10NQ3 | *VPS9A* | RRDSDASSNPVERVQ**S**ISDLEKKGAAELLKD | S337 | 0.006362 | 0.083184 | -0.241907 | V | E | R | V | Q | **S** | I | S | D | L | E | Hu et al., 2022; Hu et al., 2024 |
| A3AYP1 | *Os04g0679400* | AIFQKQVAHAPAELN**S**PRSSAAKPKNPDEIL | S18 | 0.000535 | 0.019438 | -0.359017 | P | A | E | L | N | **S** | P | R | S | S | A | Hu et al., 2022; Hu et al., 2024 |
| Q654U5 | *Os06g0247800* | FKGPNTDGGSMRQSN**S**DGALDTMARRPADPE | S722 | 0.000759 | 0.022843 | -0.400868 | M | R | Q | S | N | **S** | D | G | A | L | D | Hu et al., 2022 |
| Q67WN5 | *OVP1* | GGAWDNAKKYIEAGA**S**EHARSLGPKGSDCHK | S718 | 0.001579 | 0.036164 | -0.344055 | I | E | A | G | A | **S** | E | H | A | R | S | Hu et al., 2022 |
| Q67WN5 | *OVP1* | EGSPGAAAGKDGGAA**S**EYLIEEEEGLNEHNV | S71 | 0.373253 | 0.642851 | -0.289210 | D | G | G | A | A | **S** | E | Y | L | I | E | Hu et al., 2022 |
| Q6ZL20 | *OJ1699_E05.17* | LSIAQEAANRSATVQ**S**HEDLARKLKEEMERN | S448 | 0.001833 | 0.039365 | -0.508572 | S | A | T | V | Q | **S** | H | E | D | L | A | Hu et al., 2022 |
| A0A0P0WKD6 | *Os05g0291700* | DFSWEKLSTQLAGVA**T**QDSDEVEPKAQIATV | T426 | 0.003191 | 0.052913 | -0.320021 | L | A | G | V | A | **T** | Q | D | S | D | E | Hu et al., 2022 |

**Figure S12: Conserved SnRK1 target motifs identified in rice.** Phosphosites corresponding to proteins found in our dataset and previously reported in the literature as SnRK1 targets in rice. Columns include the UniProt protein ID, gene name, the 15-amino-acid phosphosite sequence window (centered on the phosphorylated residue), position of the phospho-acceptor (S or T) within the full-length protein, nominal p-value, adjusted p-value (p-adj.), and $log_2$ fold change ($log_2FC$) comparing *snrk1bc* to WT.
