## Supplementary material for "Integrative transcriptomic and phosphoproteomic analysis reveals key components of SnRK1 signaling network in rice": Fig. S1 - S12

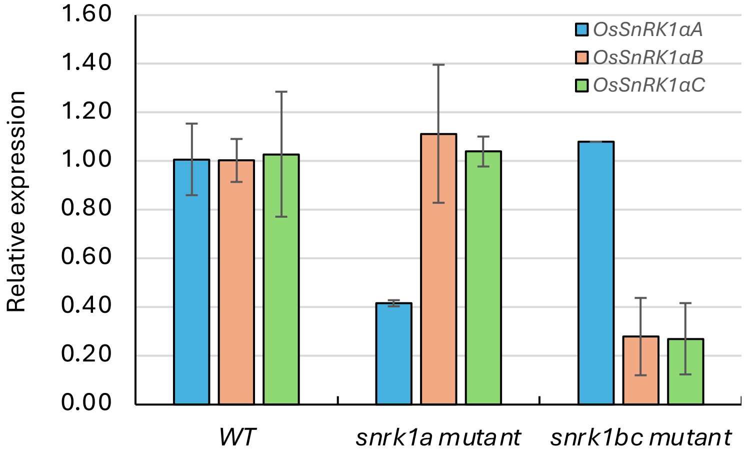


**Figure S1: Gene expression analysis of *OsSnRK1α* genes in rice *snrk1* mutants.** Gene expression of *OsSnRK1A*, *OsSnRK1B*, and *OsSnRK1C* genes determined by real-time qPCR and normalized against *OsUBQ2* gene in 9-d-old seedlings of WT, *snrk1a*, *snrk1bc* mutants grown in 14 h photoperiod on MS½ media. The graph shows expression relative to the value for gene in WT. Mean of 2 – 3 biological replicates are shown with ± standard deviation (SD).

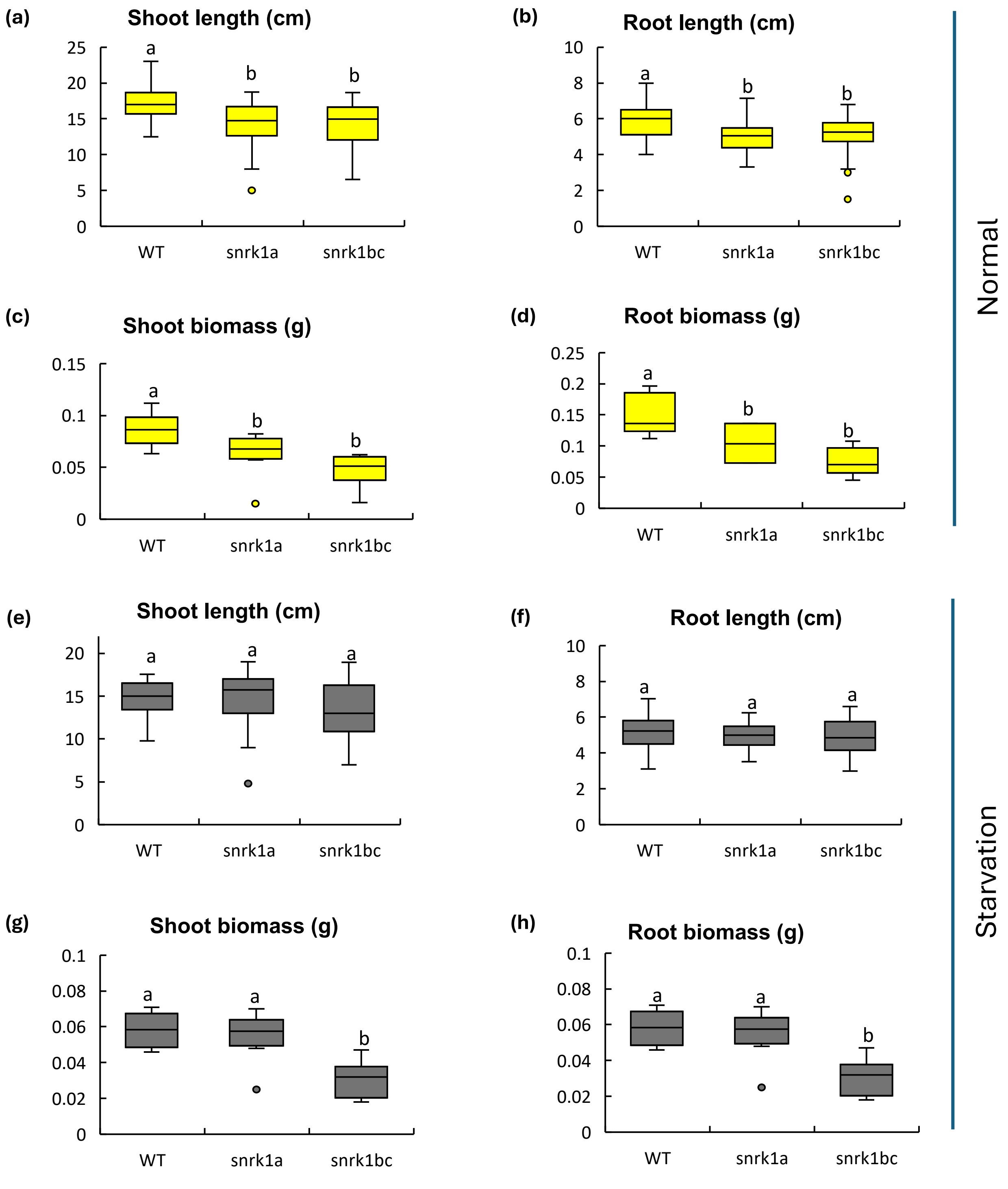


**Figure S2: Phenotypic characterization of *snrk1* mutants under normal and starvation conditions. (a–d)** Boxplots showing shoot length **(a)**, root length **(b)**, shoot biomass **(c)**, and root biomass **(d)** of 9-day-old WT, *snrk1a*, and *snrk1bc* seedlings grown under normal (14-h light/10-h dark) conditions. **(e–f)** Boxplots showing shoot length **(e)**, root length **(f)**, shoot biomass **(g)**, and root biomass **(h)** of seedlings grown for 7 days under normal condition and then exposed to 48 h of continuous darkness (starvation). Different letters indicate significant differences ($p$<0.05) determined by one-way ANOVA followed by Tukey's HSD test on data collected from 10 to 24 seedlings. Error bars represent standard deviation (SD).

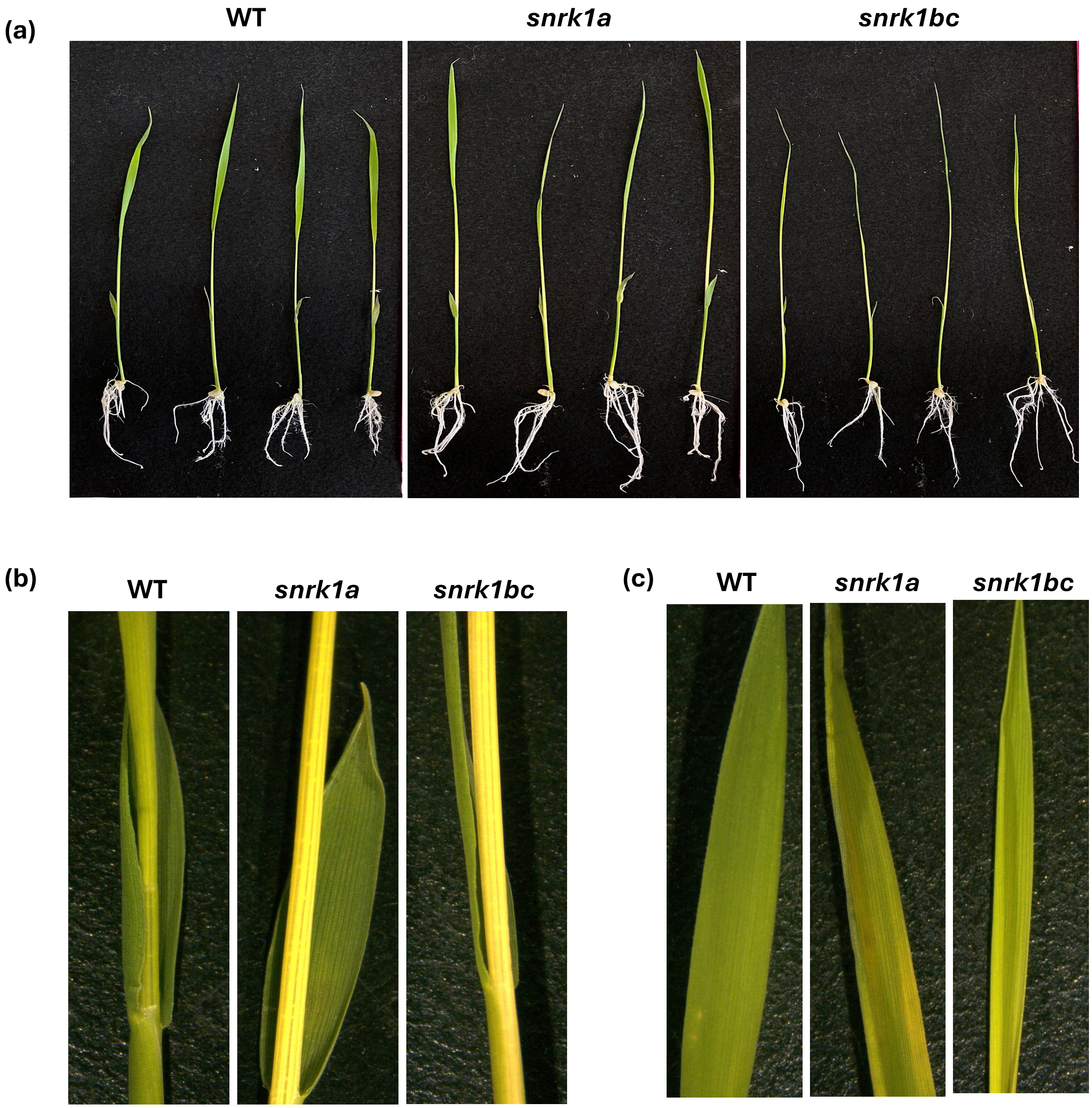


**Figure S3. Starvation-induced phenotype. (a)** Representative images of seedlings of WT, *snrk1a*, and *snrk1bc* after 48 h of continuous darkness (starvation) imposed on 7-d-old seedlings. (**b-c**) close-up of the culm at the first leaf collar (b) and fully expanded leaf (c).

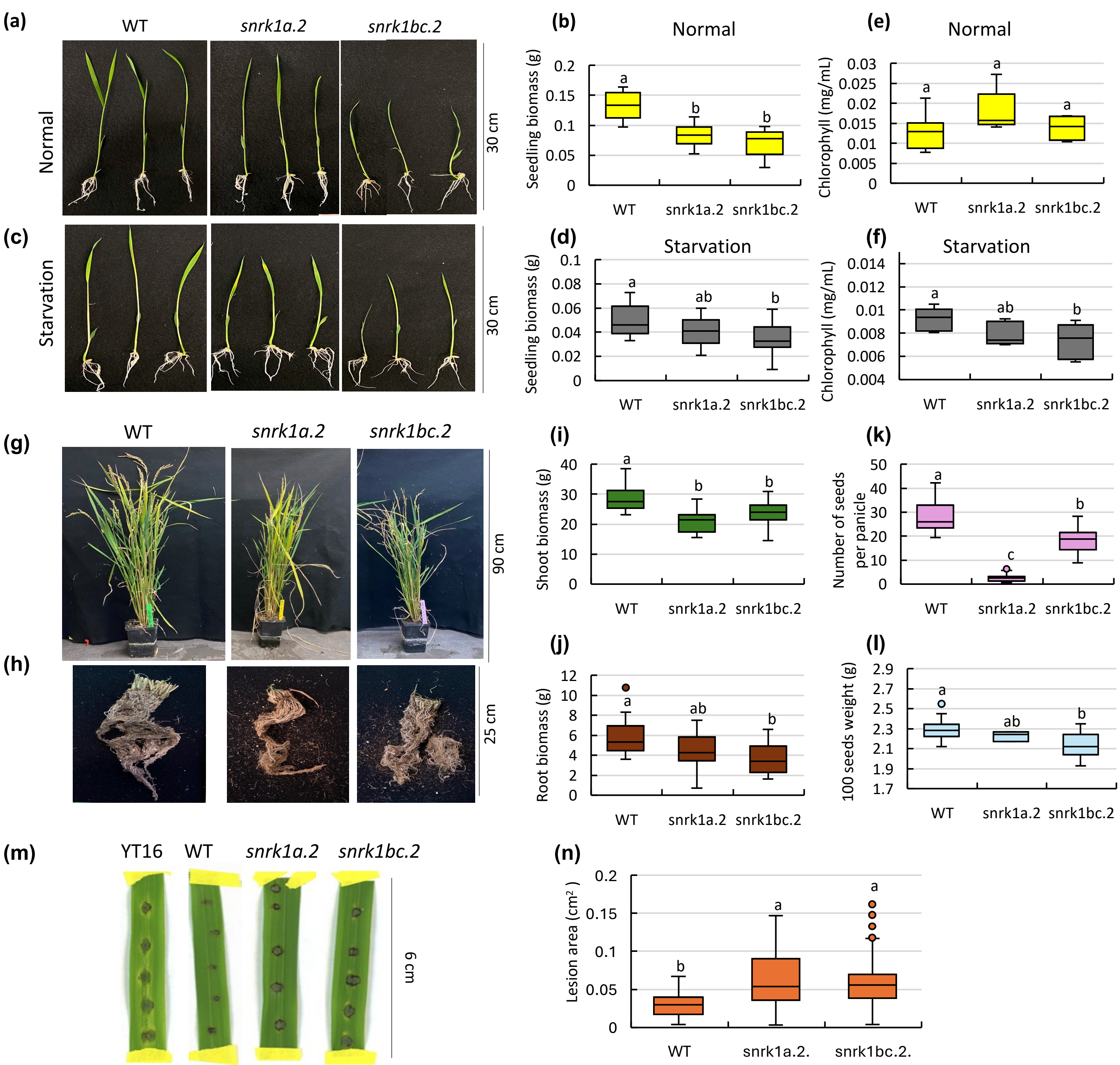


**Figure S4:** Phenotypic characterization of *snrk1a.2* and *snrk1bc.2* (second alleles of *ossnrk1aa* and *ossnrk1abc*). **(a-d)** Seedling phenotype and biomass of 9-d-old seedlings grown on half-strength MS medium under 14-h photoperiod (normal condition) or exposed to 48 h of continuous darkness after 7d of normal growth (starvation). **(e–f)** Chlorophyll content in 9-d-old seedlings under normal or starvation condition. **(g–l)** Plants at maturity in greenhouse conditions. Representative shoot and root images (g-h), shoot and root biomass (i-j), and reproductive traits (k-l). **(m-n)** Disease response of detached leaves following inoculation with *M. oryzae* strain Guy11. YT16 is a susceptible rice. Significant differences ($p$<0.05) by one-way ANOVA followed by Tukey's HSD test is shown by small betters. Error bars represent standard deviation (SD). n= 16 (a-f), n = 20 (g-l), n = 18 (m-n).

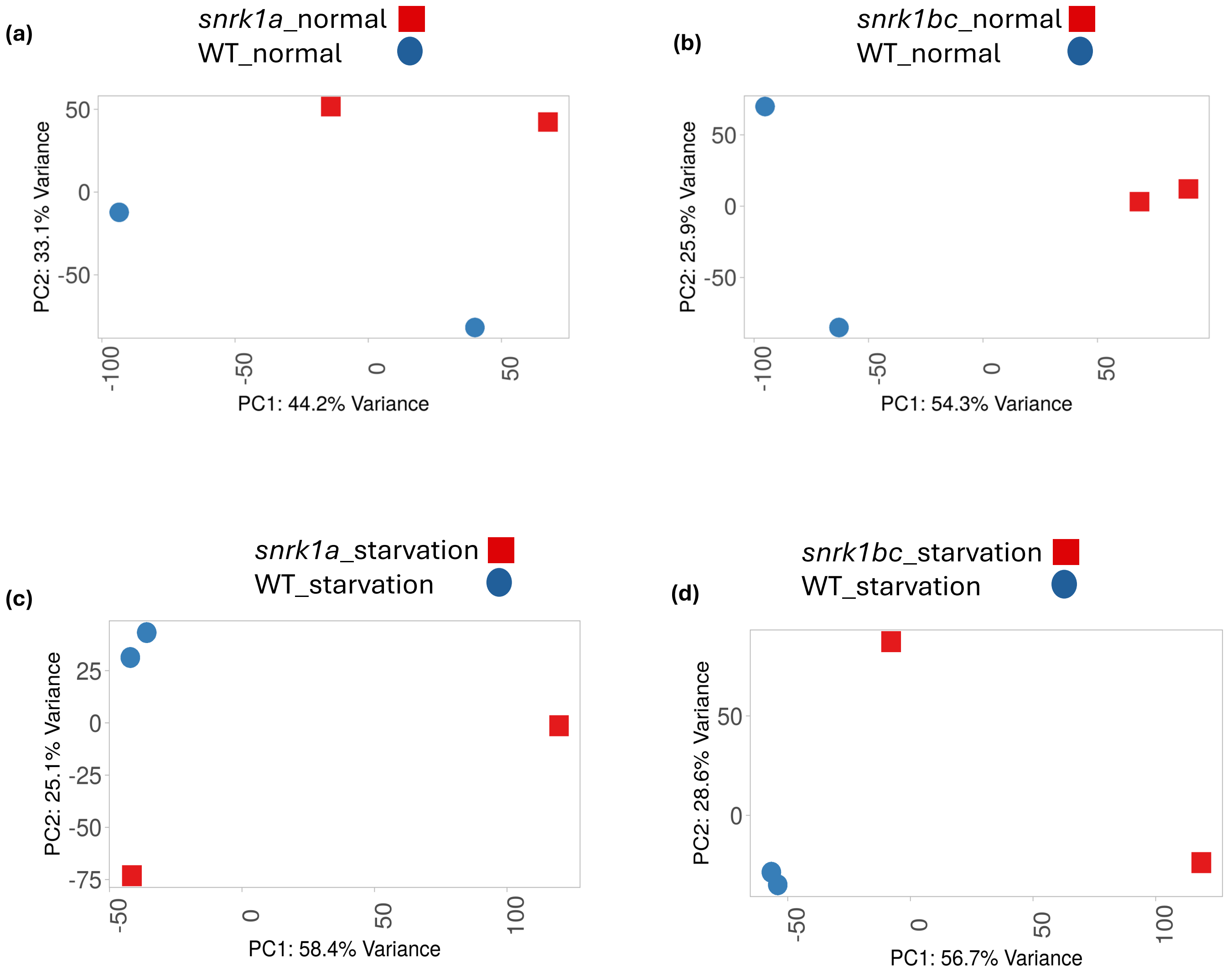


**Figure S5. Principal component analysis (PCA) of transcriptomic profiles of *snrk1* mutants and WT under normal or starvation conditions.** PCA plots of RNA-seq data (normalized gene count) from 9-d-old WT, *snrk1a*, and *snrk1bc* seedlings grown under normal condition (14-h photoperiod) or subjected to 48 h of continuous darkness after 7 days of normal growth (starvation condition). PCA plots were generated on iDEP web-based tool (Ge et al., 2018).

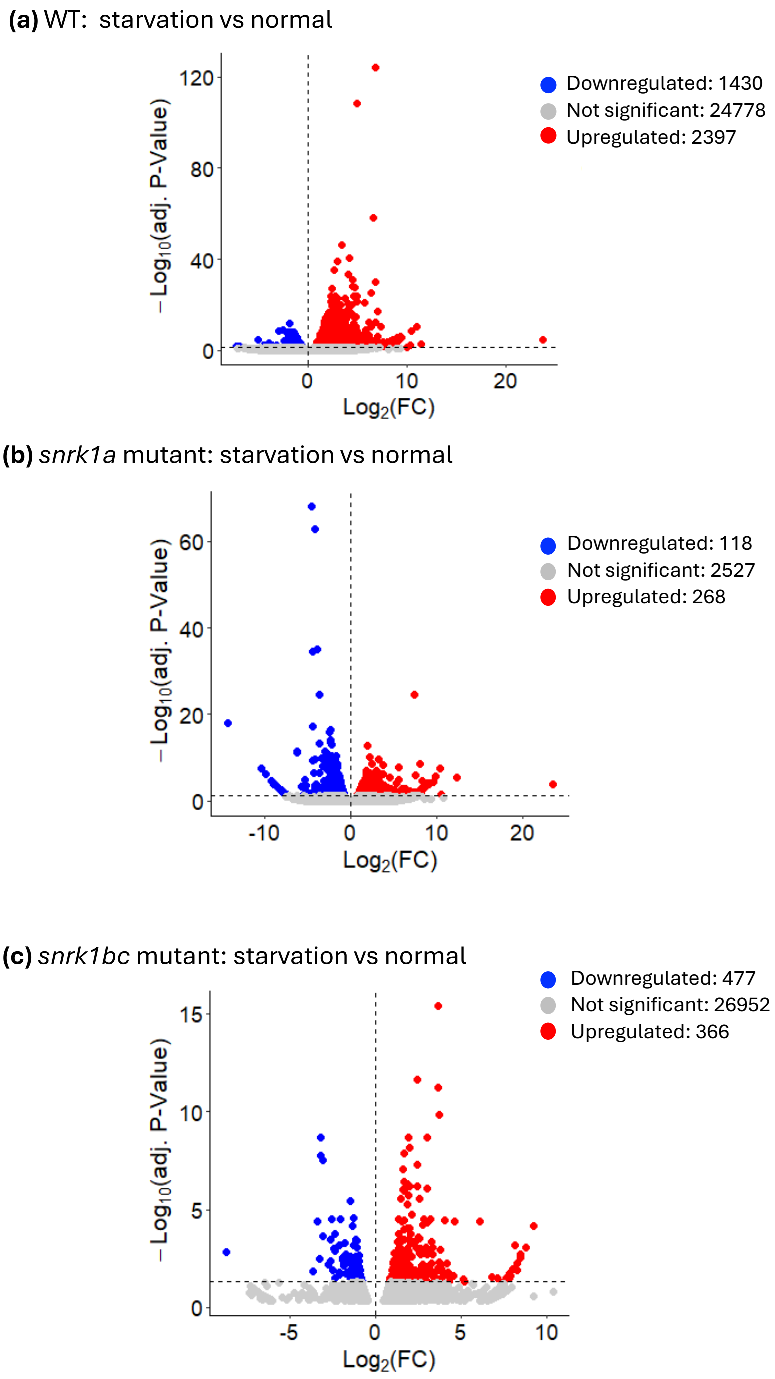


**Figure S6. Differential gene expression in WT and *snrk1* mutants under starvation compared to normal growth.** Volcano plots showing transcriptomic changes in **(a)** WT, **(b)** *snrk1a*, and **(c)** *snrk1bc* seedlings after 48 h of continuous darkness (starvation) versus those grown in normal condition. Differentially expressed genes (DEGs) were identified based on P-adj.< 0.05 and $|\log_2 FC| > 0$.

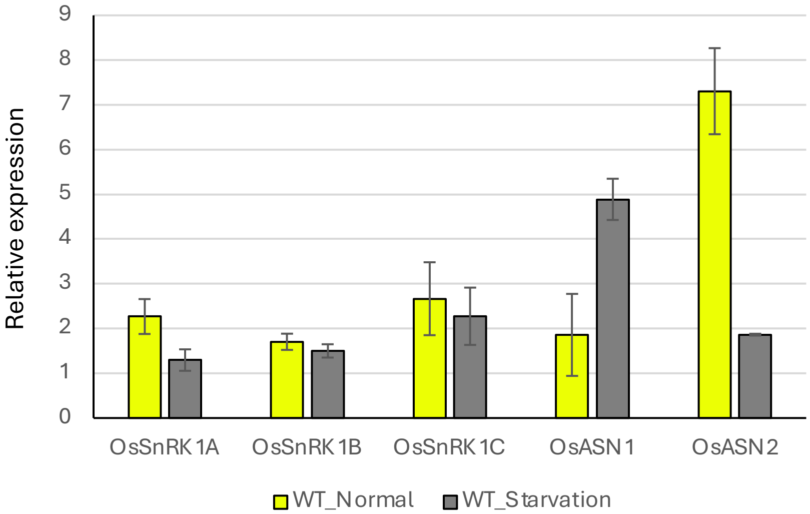


**Figure S7: Gene expression analysis of Os*SnRK1α* subunit genes and rice homologs of Arabidopsis dark-induced 6 (*DIN6*).** qPCR analysis on the rice *OsSnRK1α* paralogs (A, B, C) and the rice homologs of Arabidopsis *DARK INDUCIBLE 6/ASPARAGINE SYNTHASE 1* (*DIN6/ASN1*) (*OsASN1* and *OsASN2*) in 9-d-old seedlings of WT Kitaake seedlings grown in 14 h photoperiod (normal) or exposed to 2 days of continuous darkness after 7 days of normal growth (starvation). Gene expression was normalized against *OsUBQ2* gene (Os02g0161900). Average of 2 – 3 samples (biological replicates) was plotted with standard deviation shown as error bars.

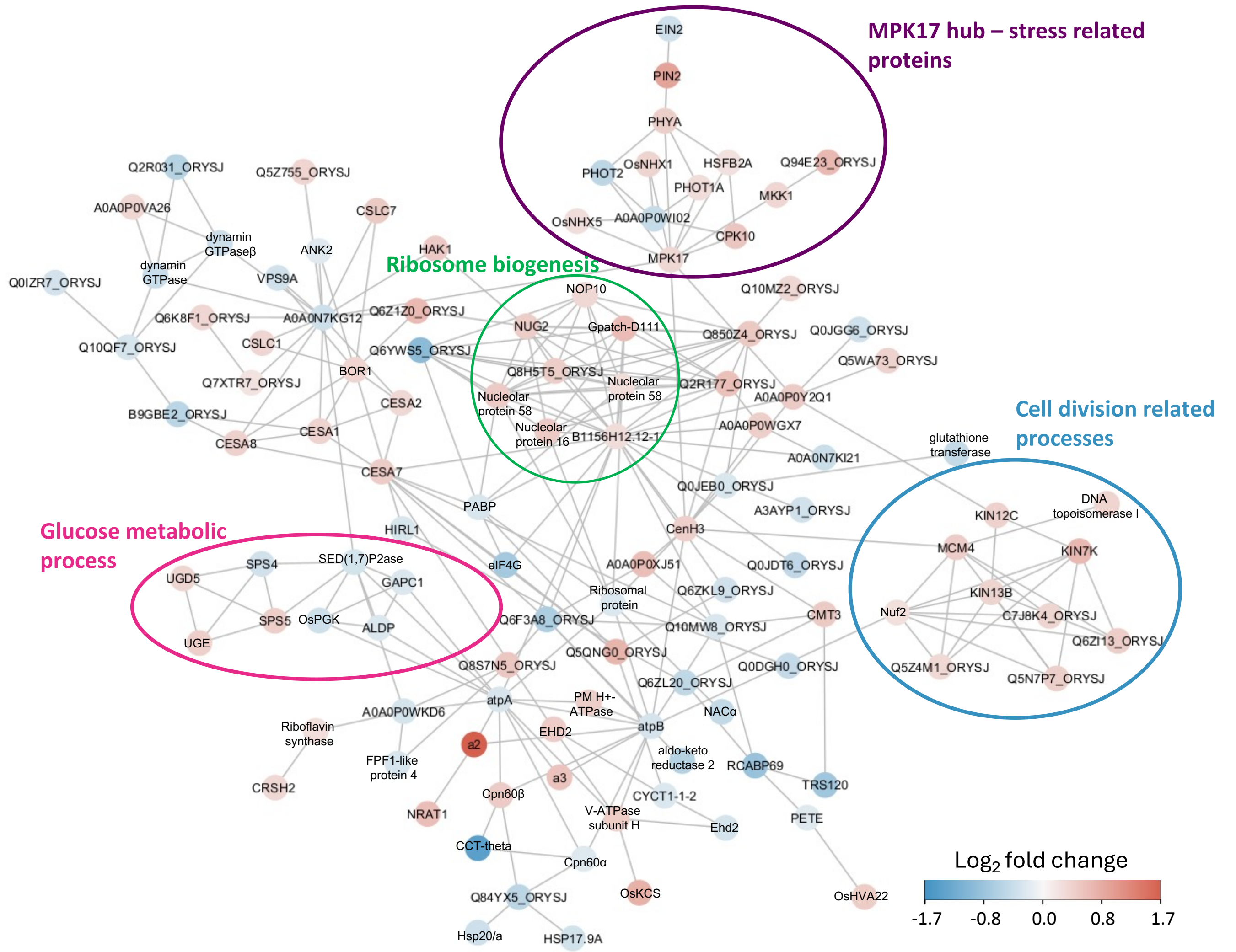


**Figure S8: Protein interaction network associated with differentially-abundant phosphosites in *snrk1bc* mutant under extended darkness.** STRING-based network analysis of significantly abundant phosphosites in *snrk1bc* seedlings under starvation (p-adj.< 0.055; $|\log_2$ fold change$| > 0$). Nodes represent proteins and edges represent known or predicted protein–protein interactions. Nodes are color-coded by $\log_2$ fold change: red indicates proteins associated with upregulated phosphosites and blue indicates proteins associated with downregulated phosphosites. Selected networks associated with gene ontology (GO) terms are circled.

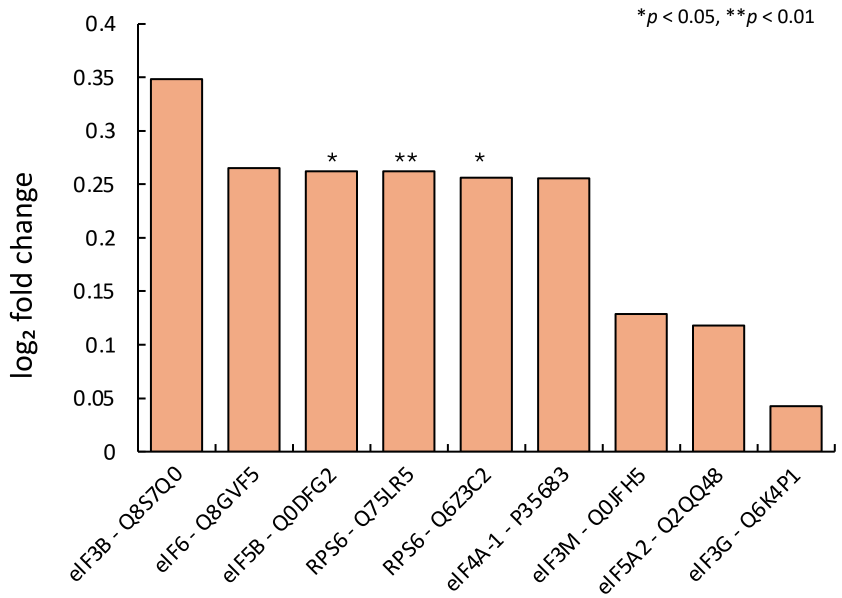


**Figure S9: Phosphorylation status of conserved TOR-S6K targets in *snrk1bc*.** Enhanced phosphorylation ($\log_2$ fold change) of ribosomal protein S6 (RPS6) and eukaryotic translation initiation factors (eIFs) in *snrk1bc* compared to WT under starvation. Significance (*p*-value) is indicated as * or **. The rest of the phosphosites show non-significant fold-change.

**(a)**

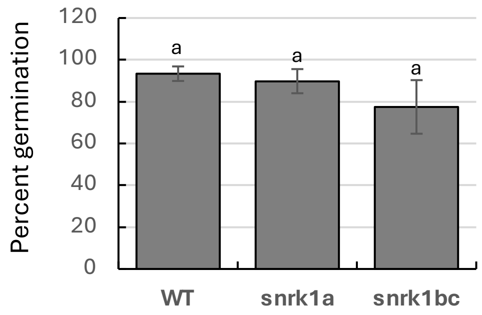


**(b)**

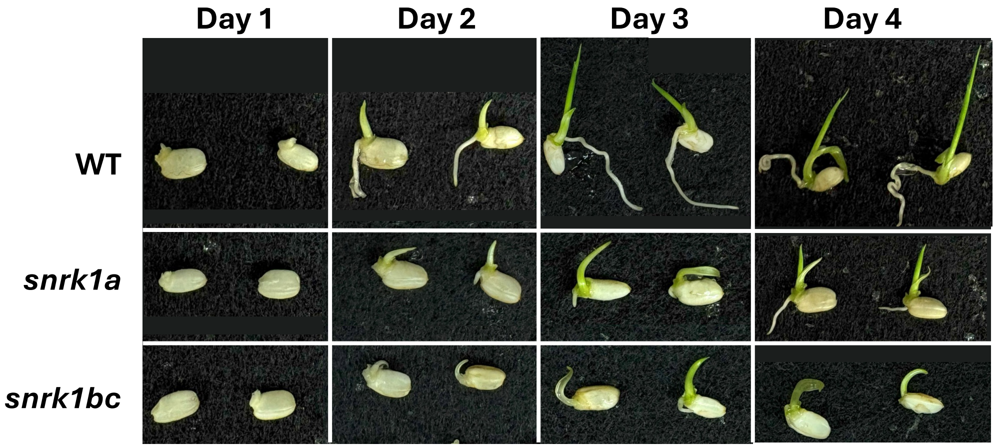


**Figure S10: Seed germination on half-strength MS media. (a)** percent germination based on number of seeds showing emergence of coleoptile after 3 days on the media at 28°C in 14 h photoperiod (n=70). **(b)** Representative images of germinating seeds of WT and *snrk1* mutants on the media. Note slower rate of shoot and root elongation in *snrk1* mutants, although percent germination is not significantly different ($p$<0.05) by one-way ANOVA followed by Tukey's HSD test.

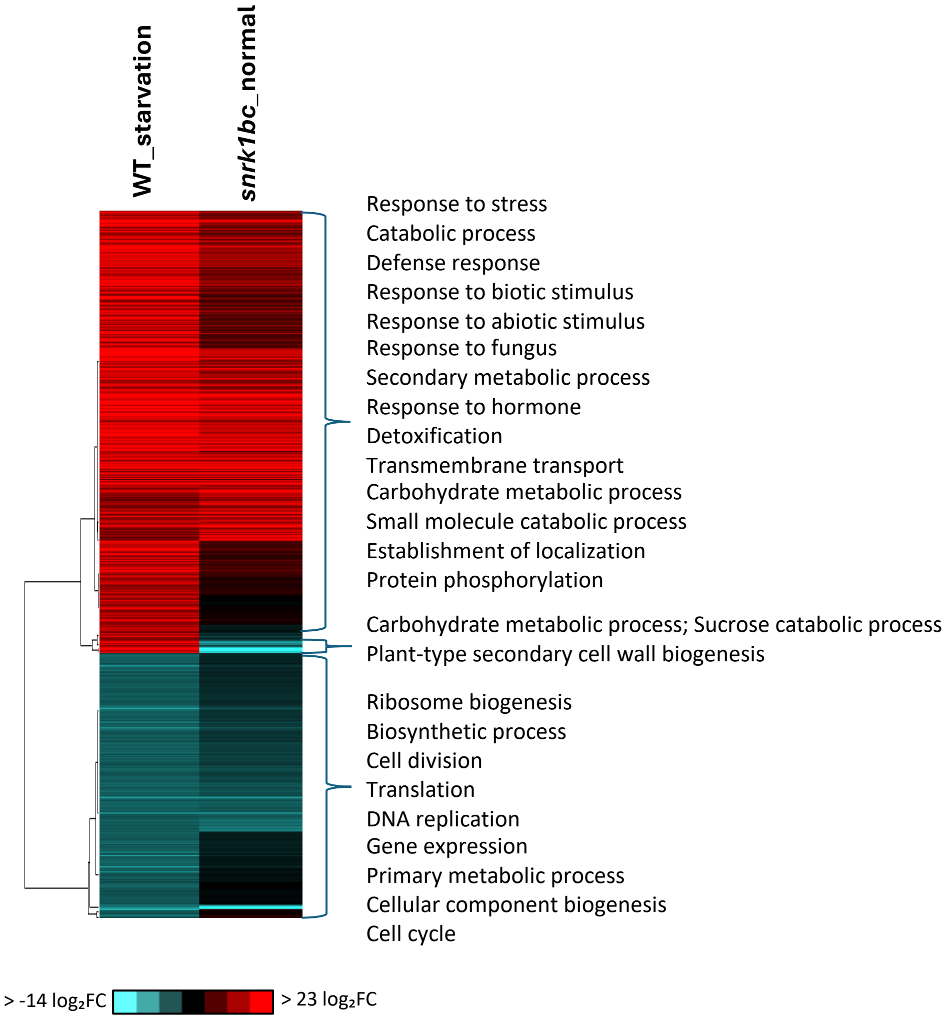


**Figure S11: Transcriptomic profile of *snrk1bc* under normal condition mirrors the transcriptomic profile of WT under starvation.** Hierarchical clustering of differentially expressed genes in WT under starvation (WT_starvation vs WT_normal) and *snrk1bc* mutant under normal condition (*snrk1bc*_normal vs WT_normal). Genes were clustered based on fold-change, with red indicating upregulation and blue indicating downregulation. Associated gene ontology (GO) terms or functional keywords are annotated.
